# Enhancer Poising Enables Pathogenic Gene Activation by Noncoding Variants

**DOI:** 10.1101/2025.06.20.660819

**Authors:** Ethan W. Hollingsworth, Zhuoxin Chen, Cindy X. Chen, Sandra H. Jacinto, Taryn A. Liu, Evgeny Z. Kvon

**Affiliations:** Department of Developmental and Cell Biology, University of California, Irvine, CA 92697, USA; Medical Scientist Training Program, University of California, Irvine School of Medicine, Irvine, CA 92697, USA

**Keywords:** enhancer variants, poised enhancer, pioneer TF, cis-regulatory elements, SNP, SNV

## Abstract

Single nucleotide variants within enhancers—non-coding DNA elements that regulate transcription—often lead to aberrant gene activation and contribute to a wide range of genetic disorders (Claussnitzer et al. 2015; Doan et al. 2016; Turner et al. 2017; Yanchus et al. 2022; Lettice et al. 2008). The mechanism by which ectopic gene activation occurs through these gain-of-function enhancer mutations remains poorly understood. Using the ZRS, a benchmark disease-associated enhancer of *Sonic hedgehog* (*Shh*), as a model, we demonstrate that poised (i.e., accessible but inactive) chromatin sensitizes *Shh* to aberrant activation in anterior limb bud, leading to polydactyly. In the anterior limb cells of wild-type mice, *Shh* is inactive, but the ZRS is accessible and marked by enhancer-associated histone modifications. We demonstrate that this poising signature explains how over 20 independent rare variants within the ZRS cause *Shh* misexpression in the same anterior limb bud cell population, resulting in similar limb malformations, despite affecting binding sites for different activators and repressors. Disabling pioneer transcription factor binding to the ZRS suppresses its poised state in anterior cells, prevents aberrant activation of the ZRS by rare variants, and fully rescues limb malformations in variant knock-in mice. A thorough examination of other disease-associated enhancers with pathogenic gain-of-function variants revealed that they are all poised in tissues with ectopic activity. We use this poising signature to predict and validate *in vivo* ectopic forebrain activity of previously uncharacterized autism-associated non-coding variants. Our findings suggest that spatial enhancer poising, likely a byproduct of development, creates a susceptibility to non-coding mutations and offers a potential mechanistic explanation for the burden of disease-associated non-coding variants.

## Introduction

Human genetics studies generate rapidly expanding lists of non-coding variants potentially causing or contributing to human diseases (Mullins et al. 2021; Zhao et al. 2021; Bellenguez et al. 2022; Trubetskoy et al. 2022; Liu et al. 2025; Halldorsson et al. 2022; Hatzikotoulas et al. 2025; S. Chen et al. 2024; Mishra et al. 2022). Most of these non-coding variants are hypothesized to affect transcriptional enhancers, abundant DNA regulatory sequences that activate gene expression in a cell-type-specific manner and in response to internal and external stimuli (Kim and Wysocka 2023; McGuire et al. 2020; Zaugg et al. 2022; Maurano et al. 2012). There is a significant gap in our understanding of how variants in enhancers alter gene expression and what impact these non-coding variants have on an organism. Among the least understood are common and rare gain-of-function enhancer variants that cause an increase in gene expression and contribute to neurodevelopmental disorders (Doan et al. 2016; Turner et al. 2017; Padhi et al. 2021; De Vas et al. 2023), cancers (Dunning et al. 2016; Bailey et al. 2016; Hua et al. 2018; Yanchus et al. 2022), and congenital malformations (Lettice et al. 2008; Hill and Lettice 2013), among other disorders (Claussnitzer et al. 2015; Stankey et al. 2024; Gupta et al. 2017). Nearly two-thirds of all experimentally studied enhancer variants linked to human disease are thought to work through an increase in enhancer activity and corresponding gene expression (**Fig. S1A and Table S1**). Consistent with this, recent large-scale massive parallel reporter assay (MPRA) studies show that 50-60% of candidate enhancer variants increase reporter gene expression (Rummel et al. 2023; Lee et al. 2025; Deng et al. 2024) (**Fig. S1B**).

Mechanistically, gain-of-function enhancer variants can disrupt a repressor binding site (Yanchus et al. 2022; Lettice et al. 2017; Vicente et al. 2015), create an activator binding site *de novo* (Fogarty et al. 2014; T. Shen et al. 2022; Downes et al. 2021; Soldner et al. 2016; Bigot et al. 2016) or increase the affinity of an existing activator binding site (Farley et al. 2015; T. Shen et al. 2022; Jindal et al. 2023; Lim et al. 2024). However, many of the affected transcriptional activators and repressors are broadly expressed during development, yet gain-of-function enhancer variants typically cause highly cell-type-specific ectopic gene expression, often in cells where the enhancer is not normally active (Claussnitzer et al. 2015; Doan et al. 2016; Turner et al. 2017; Kvon et al. 2020; Eufrásio et al. 2020; Yanchus et al. 2022). The mechanisms underlying the cell-type-specific effects of these gain-of-function enhancer variants remain elusive, even for the most well-studied genetic loci (Kvon et al. 2020; Lim et al. 2024). Identifying this missing layer of gene regulation is critical for the functional interpretation of the rapidly growing lists of disease-linked non-coding variants.

Here, we present evidence that enhancer poising by pioneer transcription factors (TFs) enables aberrant gene activation by pathogenic non-coding variants. Focusing on the limb-specific ZRS enhancer of *Shh*, which harbors 25 independent human variants linked to limb malformations, we find that chromatin poising by pioneer factor ZIC3 is required for ectopic ZRS activation in mouse embryos. In the absence of ZIC3 binding, the ZRS enhancer is no longer susceptible to gain-of-function mutations, resulting in mice that are resistant to variant-induced congenital limb malformations. We identify a similar poised signature at all other previously characterized disease-linked enhancers with gain-of-function variants. We use this poised signature to predict the cell-specific effects of previously untested human enhancer variants associated with autism. Our results indicate that bystander enhancer poising during mammalian development makes enhancers vulnerable to pathogenic non-coding mutations.

## Results

### Ectopic enhancer activity occurs in tissues with poised chromatin

To gain insights into the mechanism of ectopic gene activation by non-coding mutations, we systematically examined all previously characterized *in vivo* human developmental enhancers that contain experimentally verified gain-of-function disease-linked variants or artificial mutations. This panel consisted of 18 human enhancers containing 54 common and rare gain-of-function variants linked to autism, autoimmune disorders, obesity, cancers, limb defects, and congenital cranial dysinnervation disorders (Claussnitzer et al. 2015; Stankey et al. 2024; Tenney et al. 2023), as well as 18 artificial gain-of-function mutations (**Fig. S1A** and **Table S2**). In all cases, introducing gain-of-function variants or mutations causes ectopic reporter expression outside an enhancer’s native expression pattern in transgenic reporter assays *in vivo* or *in vitro* (**Figs. S1B-I** and **S2**). For example, a rare autism-linked G>A variant in the HS737 enhancer of *EBF3*, which is active in the neural tube, hindbrain, and midbrain, causes ectopic activity in the developing forebrain (**Fig. 1A**) (Turner et al. 2017; Hollingsworth et al. 2025). Likewise, a common A>G variant in the neural tube HS1709 enhancer of *MYC*, which confers a six-fold greater risk of low-grade glioma, causes gain of enhancer activity in the brain (**Fig. S1F**) (Yanchus et al. 2022).

**Figure 1.**
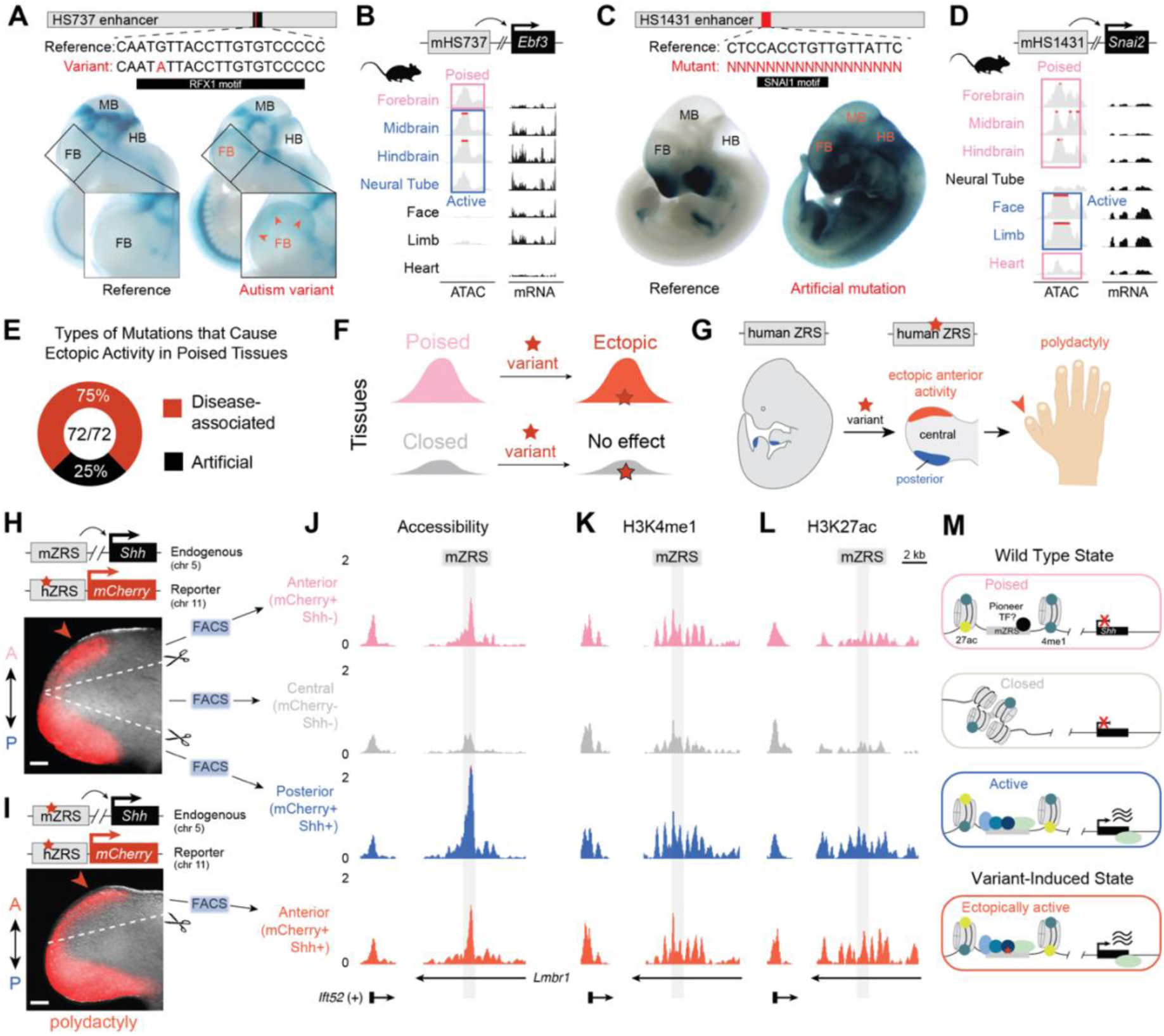
Pathogenic enhancer activation occurs in cells with a poised enhancer state. **(A)** Transgenic E11.5 whole embryos and forebrains in which a human reference HS737 enhancer allele (left) and an autism-linked HS737 variant allele (right) drive LacZ (Turner et al. 2017). Red arrows denote regions with variant-induced ectopic activity. FB, Forebrain; MB, Midbrain; HB, Hindbrain. (**B**) Genomic tracks of bulk tissue chromatin accessibility and gene expression for mHS737 (mouse homolog of HS737 sequence) and its target gene *Ebf3*, respectively, in wild-type E11.5 embryos. Blue, tissues in which a reference enhancer allele is active; Pink, tissues in which ectopic enhancer activity is observed. (**C**) Same as in (A) for the reference HS1431 enhancer allele (left) and an artificial HS1431 mutant allele (right) (Kosicki et al. 2024). FB, Forebrain; MB, Midbrain; HB, Hindbrain. (**D**) Same as in (B) for mHS1431 and its putative target gene *Snai2*. Embryo images were acquired from VISTA enhancer database (Kosicki et al. 2025; Visel et al. 2006). (**E)** Types of enhancer mutations in which ectopic activity occurs in tissues with poised chromatin. (**F**) Model where only tissues with poised chromatin are affected by a gain-of-function enhancer variant. (**G)** The schematic illustrates how human variants in the limb-specific ZRS enhancer of *Shh* lead to anterior expansion of ZRS activity and *Shh* expression, resulting in polydactyly. **(H, I)** Fluorescent images of E11.5 hindlimbs from a wild-type mouse containing a 404G>A variant hZRS reporter transgene driving *mCherry* (mZRS^WT^; hZRS^404G>A^-mCH) (H) and a 404G>A variant knock-in mouse with a 404G>A variant hZRS reporter transgene driving *mCherry* (mZRS^404G>A^; hZRS^404G>A^-mCH) (I). A, anterior; P, posterior. Scale bars, 500 μm. **(J-L)** Genomic tracks across FAC-sorted limb bud cell populations for levels of chromatin accessibility (J, ATAC-seq), H3K4me1 (K, CUT&Tag), and H3K27ac (L, CUT&Tag), marks at the endogenous mZRS locus (highlighted in gray). *Ift52* is shown as a positive control across cell populations. (**M**) Interpretation of observed chromatin states in different limb bud cell populations with and without a variant present.

To identify features that could correlate with or predict cell types of ectopic enhancer activity, we systematically compared normal and variant enhancer activity patterns of these 18 human enhancer regions with their respective tissue- and cell-specific chromatin states, using publicly available epigenomic datasets for both mouse and human. We assessed enhancer-associated histone modifications from ENCODE, DNA accessibility (measured by assay for transposase-accessible chromatin with sequencing (ATAC-seq) or single-nucleus ATAC-seq), and CpG methylation (**Fig. 1B** and **Figs. S1 and S2**). Consistent with their active state, all 18 enhancers are accessible and marked by active enhancer-associated histone modifications in human and mouse tissues in which they are active (**Figs. S1 and S2**). For example, the HS737 human enhancer of *EBF3* is active in the neural tube, hindbrain, and midbrain of mouse embryonic day 11.5 (E11.5) embryos and it is marked by enhancer-associated H3K27ac and H3K4me1 and is accessible in these tissues in mouse and stage-matched human fetal brain **(Figs. 1A, B and S1C)**. By contrast at E11.5, HS737 is closed and lacks the H3K27ac mark in the limb, face and heart where it is not active. Interestingly, the HS737 enhancer is accessible and marked by H3K27ac and H3K4me1 marks in the mouse forebrain, even though it is not active and *Ebf3* is not expressed in the forebrain at E11.5 (**Figs. 1B and S1C**). The G>A variant allele becomes active in the forebrain suggesting that the HS737 enhancer is poised for ectopic activation in the forebrain (**Fig. 1A**). Single-nucleus ATAC-seq from the early fetal human brain at the corresponding stage (Mannens et al. 2023) confirmed that the HS737 enhancer is likewise accessible in radial glia (RG) and neurons of the forebrain (**Fig. S1C**).

We observed a similar poised signature in all 18 enhancers examined that had tissue-matched ATAC-seq data in mouse or human; gain-of-function variants or synthetic mutations cause ectopic activity in tissues where the enhancer is accessible (**Figs. 1E, S1D-I, and S2**). In the most extreme case observed, the face/limb HS1431 enhancer of the *Snai2* gene is open in all tissues of the embryo (**Figs. 1C, D and S2E**). Consistent with this broad accessibility pattern, a synthetic gain-of-function mutation that removed a SNAI1 repressor binding site causes broad ectopic activity of the mutated HS1431 enhancer across all tissues of E11.5 embryos (**Fig. 1C, D**) (Kosicki et al. 2024). Taken together, this analysis reveals that the tissues and cell types affected by gain-of-function enhancer mutations are correlated with chromatin accessibility in both mice and humans. This correlation led us to hypothesize that having poised chromatin at a particular enhancer locus may sensitize it to ectopic gene activation (**Fig. 1F**).

### ZRS is poised in anterior limb bud cells

To further explore the role of enhancer poising in pathogenic gene activation, we focused on one of the most well-studied human enhancers, the zone of polarizing activity (ZPA) regulatory sequence (ZRS, also known as MFCS1) (Lettice et al. 2003). The ZRS is a limb-specific enhancer of *Shh* and is active in the posterior mesenchyme region of developing fore- and hindlimb buds called the ZPA. In humans, 25 independent single-nucleotide variants within the ZRS enhancer are associated with limb malformations, most commonly preaxial polydactyly the (Lettice et al. 2003; VanderMeer and Ahituv 2011; Hill and Lettice 2013; Kvon et al. 2020). ZRS mutations implicated in polydactyly cause expansion of *Shh* expression from the normal posterior ZPA domain into the anterior domain of the developing forelimb and hindlimb, resulting in extra digits (**Fig. 1G**). Given the high density of patient variants and clear phenotypic readout, the ZRS-*Shh* locus provides an advantageous model for determining the mechanisms underlying ectopic gene expression by non-coding variants. Based on the presence of the poised signature at other disease-associated enhancers (**Fig. S1C-H**), we hypothesized that the ZRS enhancer is also poised in a population of anterior limb bud cells that are prone to ectopic *Shh* expression (**Fig. 1G**).

To test this hypothesis, we sought to determine the chromatin state of the ZRS locus in posterior ZPA and anterior populations of mesenchymal limb bud cells of wild-type mice. To visualize and label these cell populations, we used a previously published transgenic mouse line in which a 404G>A ‘Cuban’ variant allele of the human ZRS (hZRS^404G>A^) drives *mCherry* in ZPA and anterior limb mesenchyme (Hollingsworth et al. 2025) (**Fig. 1H**). We dissected the anterior and posterior domains of day E11.5 hindlimb buds from hZRS^404G>A^-*mCH* transgenic mice and FAC-sorted pure populations of mCherry+ mesenchyme cells. As a negative control population, we used mCherry-cells from central hindlimb buds, which never express *Shh* (mCherry-/*Shh*-; **Figs. 1H and S3A**). We performed ATAC-seq to map accessible chromatin and CUT&Tag to profile enhancer-associated H3K4me1 and H3K27ac histone marks in anterior, central and posterior cell populations (**Fig. 1J-M**). Importantly, ATAC-seq and CUT&Tag reads from the transgenic human (hZRS) and endogenous mouse (mZRS) enhancers can be distinguished by substitutions despite the high level of conservation of the ZRS sequence (**Fig. S3B** and **Methods**).

In posterior limb bud cells (mCherry+/*Shh*+), the endogenous mZRS enhancer was highly accessible and occupied by high levels of H3K4me1 and H3K27ac, consistent with its active state (**Fig. 1J-M**). Consistent with our prediction, in anterior mCherry-positive limb bud cells of normal mice, the endogenous mZRS enhancer is also accessible, marked by H3K4me1 and H3K27ac at levels comparable to posterior limb bud cells and substantially higher than in control central limb bud cells (mCherry-/*Shh*-) (**Figs. 1J-M**). We confirmed the presence of this poised signature at the ZRS in a cluster of anterior limb bud cells using previously published single-nucleus ATAC-seq data from E11.5 hindlimb (**Fig. S1I**) (Bower et al. 2024).

To determine how the introduction of a gain-of-function mutation changes the chromatin state of the poised ZRS enhancer, we created a variant knock-in mouse line by introducing a human 404G>A variant into the endogenous mouse ZRS enhancer (mZRS^404G>A^). The resulting knock-in mice had extra digits on both forelimbs and hindlimbs, as well as a truncated tibia, faithfully reproducing phenotypes found in human patients with a 404G>A variant (Lettice et al. 2008; Wieczorek et al. 2010) (**Fig. S4A, B**). We next dissected the anterior domains of day E11.5 hindlimb buds from hZRS^404G>A^-*mCH* transgenic mice on this new variant knock-in background (mZRS^404G>A^) and FAC-sorted pure populations of mCherry+ mesenchyme cells followed by epigenomic profiling (**Fig. 1I**). In anterior limb bud cells that aberrantly express *Shh* (mCherry+/*Shh*+), the mZRS^404G>A^ allele displayed active chromatin signatures (accessible, and high levels of both H3K4me1 and H3K27ac), very similar to those observed in posterior limb bud (**Fig. 1J-N**).

Together, these data demonstrate that introducing a pathogenic variant into the endogenous mZRS enhancer converts it from a poised state (accessible, high H3K4me1 and low H3K27ac) into an active enhancer state characterized by increased H3K27 acetylation, which results in aberrant *Shh* expression and limb malformations (**Fig. 1N**).

### Independent ZRS variants cause highly overlapping ectopic activities

To explore whether ZRS enhancer poising in anterior limb bud cells can explain how other gain-of-function mutations in the ZRS cause similar limb malformations, we systematically compared the activities of independent pathogenic human ZRS variants. We have previously shown that the 404G>A variant, which disrupts a putative repressor binding site (Amano et al. 2017; Lettice et al. 2017), and another independent pathogenic 446T>A variant, which increases affinity of an HnRNP K activator binding site (Y. Chen et al. 2023), both cause highly overlapping ectopic expression in the same population of anterior limb cells (Hollingsworth et al. 2025). This led us to hypothesize that other gain-of-function variants affecting the ZRS, regardless of their location or type of affected TF (activator or repressor), would cause ectopic activity in the same population of anterior cells because they are primed for ectopic ZRS activation (**Fig. 2A**).

**Figure 2.**
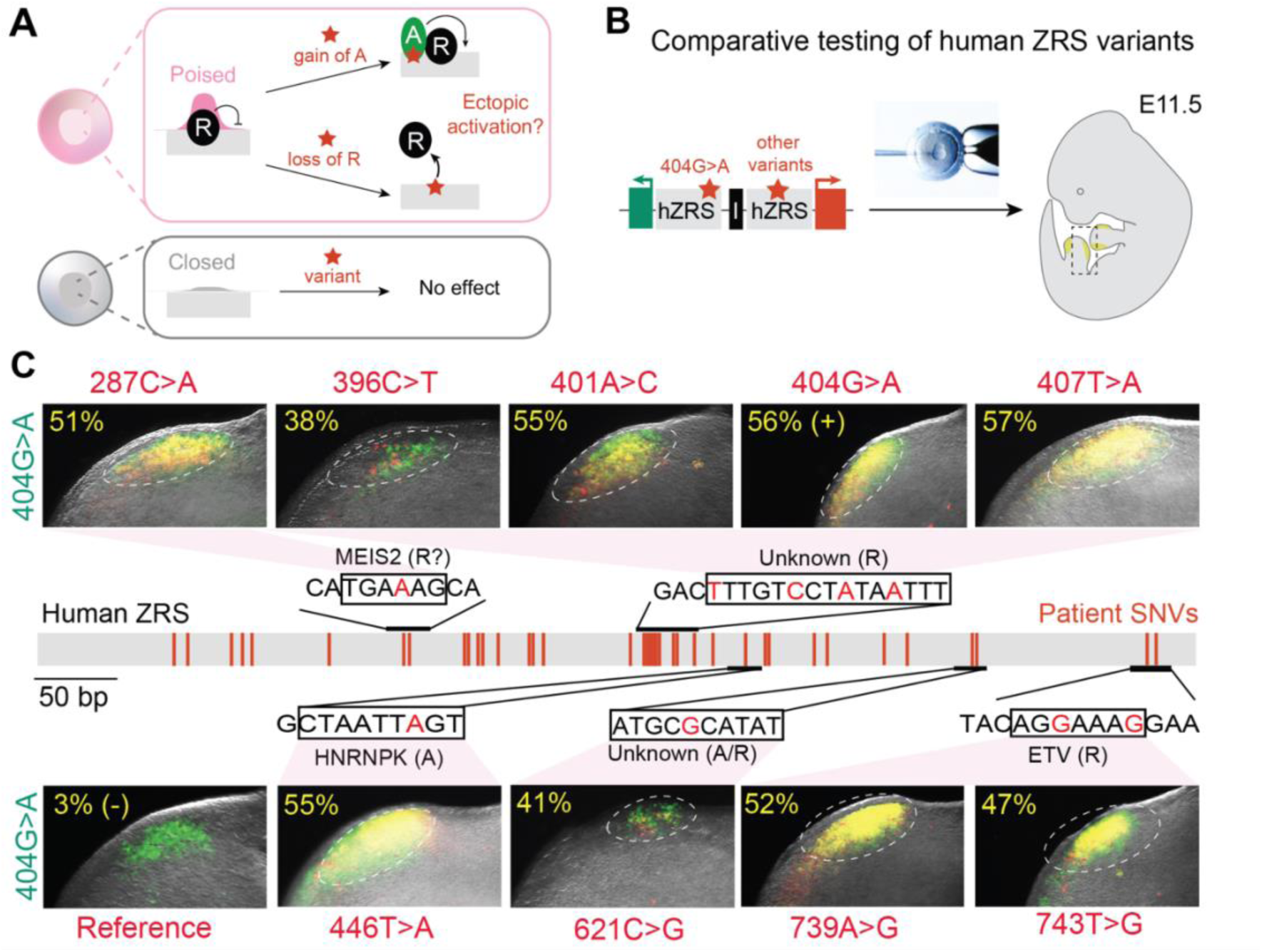
Ectopic ZRS enhancer activity driven by independent gain-of-function variants. **(A)** A model in which enhancer poising, rather than activator (A) or repressor (R) TFs, determines cell-type-specific ectopic activity. **(B)** Dual-enSERT-based strategy for comparative testing of human ZRS variants *in vivo*. **(C)** Representative fluorescent images of the anterior domains of transgenic mouse hindlimb buds in which a hZRS^404G>A^ drives *eGFP* (green channel) and different human ZRS variant alleles drive *mCherry* (red channel). Font colors of variants denote fluorophore. Yellow percentages represent FACS-based overlap in mCherry and eGFP fluorescence (**Methods**). Putative transcription factor (TF) motifs disrupted by patient variants are shown.

To test this hypothesis, we used our recently developed site-specific dual enhancer-reporter transgene system (dual-enSERT) to visualize and quantitatively compare the *in vivo* activities of different human ZRS variant alleles in the same mouse (**Fig. 2B**) (Hollingsworth et al. 2025). We created ten dual-enSERT transgenic constructs each containing a different hZRS variant allele driving *mCherry* on one side and the pathogenic human ZRS variant 404G>A allele (hZRS^404G>A^) driving *eGFP* on the other side (**Fig. 2B**). The ten different ZRS alleles we selected included the reference allele (negative control), 404G>A (positive control), 287C>A, 401A>C, 404G>A, 407T>A (all disrupt putative repressor binding), 446T>A (increases binding affinity of an activator) and a 621C>G variant of unknown etiology (**Fig. 2C**). Together, these variants (hZRS^var^) affect binding sites for at least five different TFs. We injected these constructs into mouse zygotes and examined reporter activity in transgenic F0 embryos eleven days later (see **Methods**). We observed a substantial overlap between eGFP and mCherry fluorescence in anterior limb buds of hZRS^var^-*mCherry*/hZRS^404G>A^-*eGFP* transgenic mice for all nine variant alleles but not the reference allele (**Fig. 2C**). We dissected the anterior hindlimb buds from all ten transgenic mouse lines and confirmed with FACS that the overlap in ectopic activity in the anterior limb bud cells was visually and quantitatively indistinguishable from the overlap observed in hZRS^404G>A^-*mCherry*/hZRS^404G>A^-*eGFP* transgenic embryos in which eGFP and mCherry were driven by the same hZRS^404G>A^ variant allele (All variants, *P* = ns; **Figs. 2C and S5**). Taken together, these data demonstrate that despite affecting binding sites for different activators and repressors, human variants in the ZRS cause ectopic expression in the same population of anterior limb cells in which the ZRS is poised.

### ZIC3 binding is required for anterior ZRS poising

To identify regulators of ZRS enhancer poising in anterior limb bud cells, we performed RNA-sequencing on anterior (mCherry+/*Shh*-) and central (mCherry-/*Shh*-) hindlimb bud cells from wild-type mZRS^WT^; hZRS^404G>A^-*mCH* mice followed by differential gene expression analysis (**Fig. 3A**). This analysis revealed sixteen upregulated TFs in anterior limb bud cells (**Figs. 3A**, **S6A**, and **Table S3**; log_2_-fold change > 2; adjusted *P*-value < 0.05). The most significantly upregulated gene in anterior cells was zinc finger TF *Zic Family Member 3* (*Zic3*) (Fold change = 3.14, adjusted *P*-value = 6.12 x 10^-194^). ZIC3 and its fly homolog Odd-paired act as pioneer factors to increase accessibility of enhancers in mammalian gastrulation and early fly embryogenesis (Grand et al. 2024; Soluri et al. 2020; Koromila et al. 2020) via recruitment of the SWI/SNF chromatin remodelling complex to chromatin (Hossain et al. 2024). *Zic3* is expressed in the anterior limb mesenchyme, where it is hypothesized to play a role in specifying digit number by regulating Hedgehog signaling (S. Li et al. 2025; Quinn, Haaning, and Ware 2012) (**Fig. 3B**). However, its role in the transcriptional regulation of limb developmental genes is unknown. We scanned the ZRS sequence for candidate ZIC3 binding sites and identified a highly-conserved CCTGCTG motif with a perfect match to the ZIC2 and ZIC3 consensus binding site (**Fig. 3B**). We next assessed ZIC2 and ZIC3 binding at the ZRS using CUT&TAG (ZIC2) and CUT&RUN (ZIC3) experiments in anterior and posterior E11.5 limb bud cells. While we observed strong ZIC2 binding at selected loci genome-wide, we found no evidence for ZIC2 binding at the ZRS region in anterior or posterior limb bud cells at E11.5 (**Figs. 3C** and **S6B, C**). By contrast, ZIC3 is strongly bound to the ZRS in anterior limb bud cells (**Fig. 3D**).

**Figure 3.**
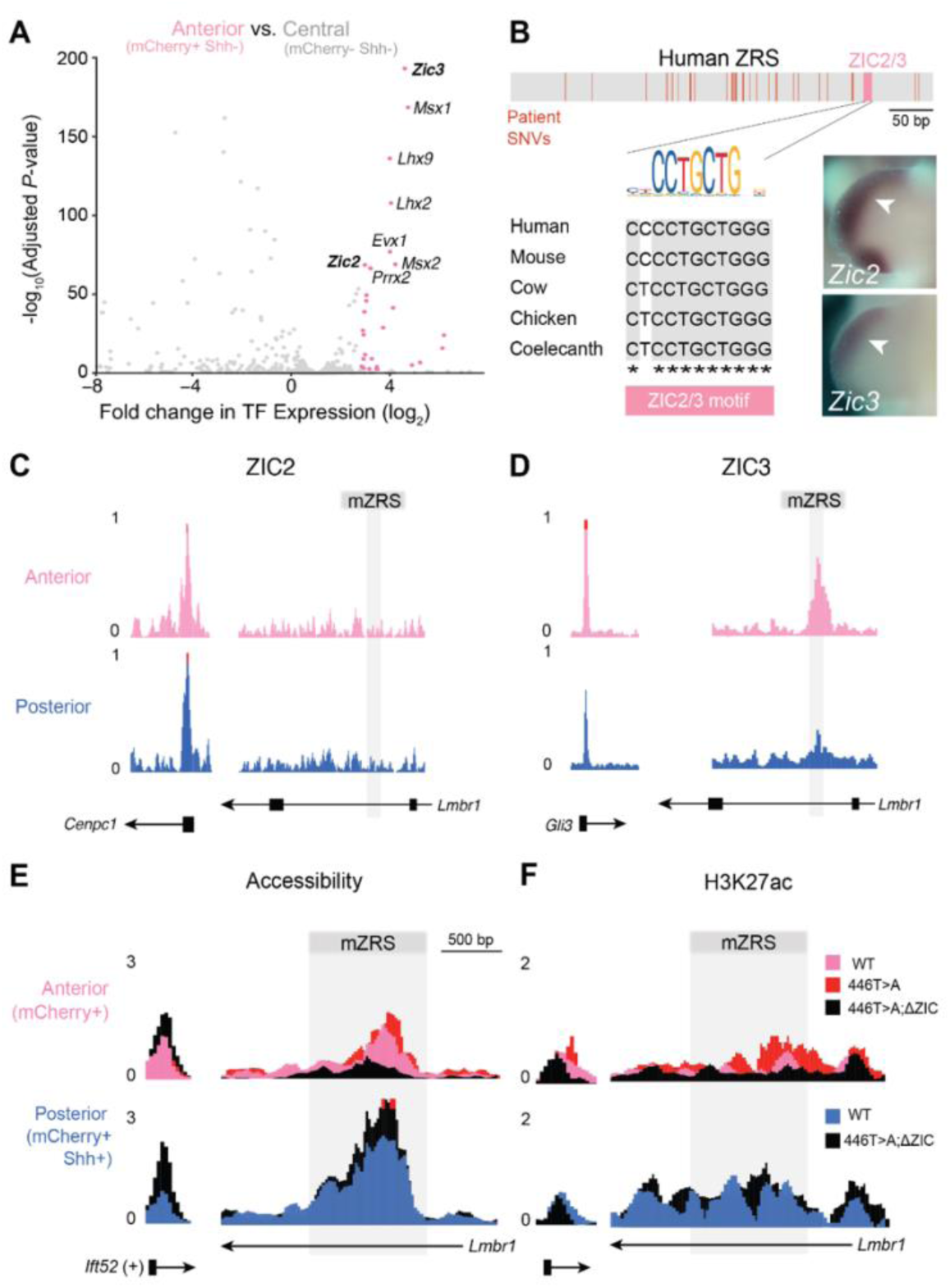
ZIC3 binding is required for anterior poising at the ZRS. (**A**) Volcano plot of TF expression between anterior (mCherry+/Shh-) and central (mCherry-/Shh-) limb bud cells from mZRS^WT^;hZRS^404G>A^-*mCH* hindlimb buds. Significantly upregulated TFs (adjusted P-value < 0.05 and log_2_(fold-change) > + 2) are colored in pink. (**B**) Location and evolutionary conservation of putative binding motif for Zic Family Member TFs 2 and 3 (ZIC2/3) within the human ZRS enhancer (pink). Patient SNVs in red. *Zic2* and *Zic3* mRNA whole mount in situ hybridization in E11.5 limb buds (bottom right). Images are reproduced from the Embryos database (http://embrys.jp). **(C-D)** ZIC2 CUT&Tag and ZIC3 CUT&RUN occupancy in anterior and posterior E11.5 hindlimb buds at the mouse ZRS and control loci. **(E-F)** Genomic tracks of chromatin accessibility (E, ATAC-seq), and H3K27ac (F, CUT&Tag) signal at the mZRS locus in FAC-sorted anterior cells (top) from mZRS^WT^;hZRS^446T>A^-*mCH* (pink), mZRS^446T>A^;hZRS^446T>A^-*mCH* (red) and mZRS^446>T>AΔZIC^;hZRS^446T>A^-*mCH* (black) mice and posterior cells (bottom) from mZRS^WT^;hZRS^446T>A^-*mCH* (blue) and mZRS^446>T>AΔZIC^;hZRS^446T>A^-*mCH* (black) mice. *Ift52* is shown as a positive control between populations and genotypes.

To determine the extent to which ZIC3 binding is required for anterior ZRS poising we created a knock-in mouse line in which we replaced endogenous mZRS with a variant pathogenic mZRS allele along with a mutagenized ZIC2/3 motif (mZRS^446T>A;ΔZIC^; **Fig. S7A, B**). To assess chromatin state at the ZRS in the absence of ZIC3 binding, we generated mZRS^446T>A;ΔZIC/446T>A;ΔZIC^; hZRS^446T>A^-*mCH* mice. We isolated anterior mCherry+/Shh-, central mCherry-/Shh-, and posterior mCherry+/Shh+ cells from dissected E11.5 hindlimbs and performed ATAC-seq and CUT&Tag for H3K27ac mark. Remarkably, removal of the ZIC3 binding site substantially reduced mZRS chromatin accessibility and H3K27ac occupancy in anterior limb bud cells, to the levels comparable to central limb bud cells (**Figs. 3E, F** and **S6D, E**). In comparison, the chromatin accessibility and the occupancy of H3K27ac mark were not affected by ZIC3 motif mutagenesis in posterior limb bud cells. (**Fig. 3E-G**). Together, these data show that ZIC3 binding is required for chromatin opening at the ZRS in anterior limb bud cells.

### Anterior poising is required for aberrant enhancer activation and limb malformations

To test whether the anterior poised signature in the ZRS is required for ectopic enhancer activity caused by gain-of-function mutations *in vivo*, we performed comparative transgenic reporter experiments. As a point of comparison, we used transgenic mouse lines in which 404G>A (decreased repressor binding) and 446T>A (increased activator binding) variant ZRS alleles drove ectopic *mCherry* expression in anterior limb bud cells and a line in which *mCherry* is driven by a reference ZRS allele (**Fig. 4A**). We mutagenized a single pioneer factor ZIC2/3 motif in both variant ZRS alleles, placed them upstream of *mCherry*, and injected the resulting construct into fertilized mouse embryos. Live imaging revealed a complete loss of mCherry fluorescence in anterior limb buds of hZRS^404G>A;ΔZIC^-*mCH* and hZRS^446G>A;ΔZIC^-*mCH* transgenic mice in fore- and hindlimbs (hZRS^404G>A;ΔZIC^: Forelimb, *P* = 2.51E-05; Hindlimb, *P* = 7.46E-06. hZRS^446G>A;ΔZIC^: Forelimb, *P* = 1.40E-06; Hindlimb, *P* = 4.05E-06) while posterior mCherry fluorescence was not affected (*P* = ns; **Figs. 4A, B** and **S7D, E**). In fact, the enhancer activity patterns of variant alleles lacking the ZIC2/3 motif were indistinguishable from enhancer activity driven by a reference ZRS allele (*P* = ns for all) (**Figs. 4A, B and S7D**). Together, these results indicate that the ZIC2/3 motif is not required for normal posterior ZRS expression domain but is necessary for ectopic anterior activity caused by multiple, mechanistically distinct ZRS mutations.

**Figure 4.**
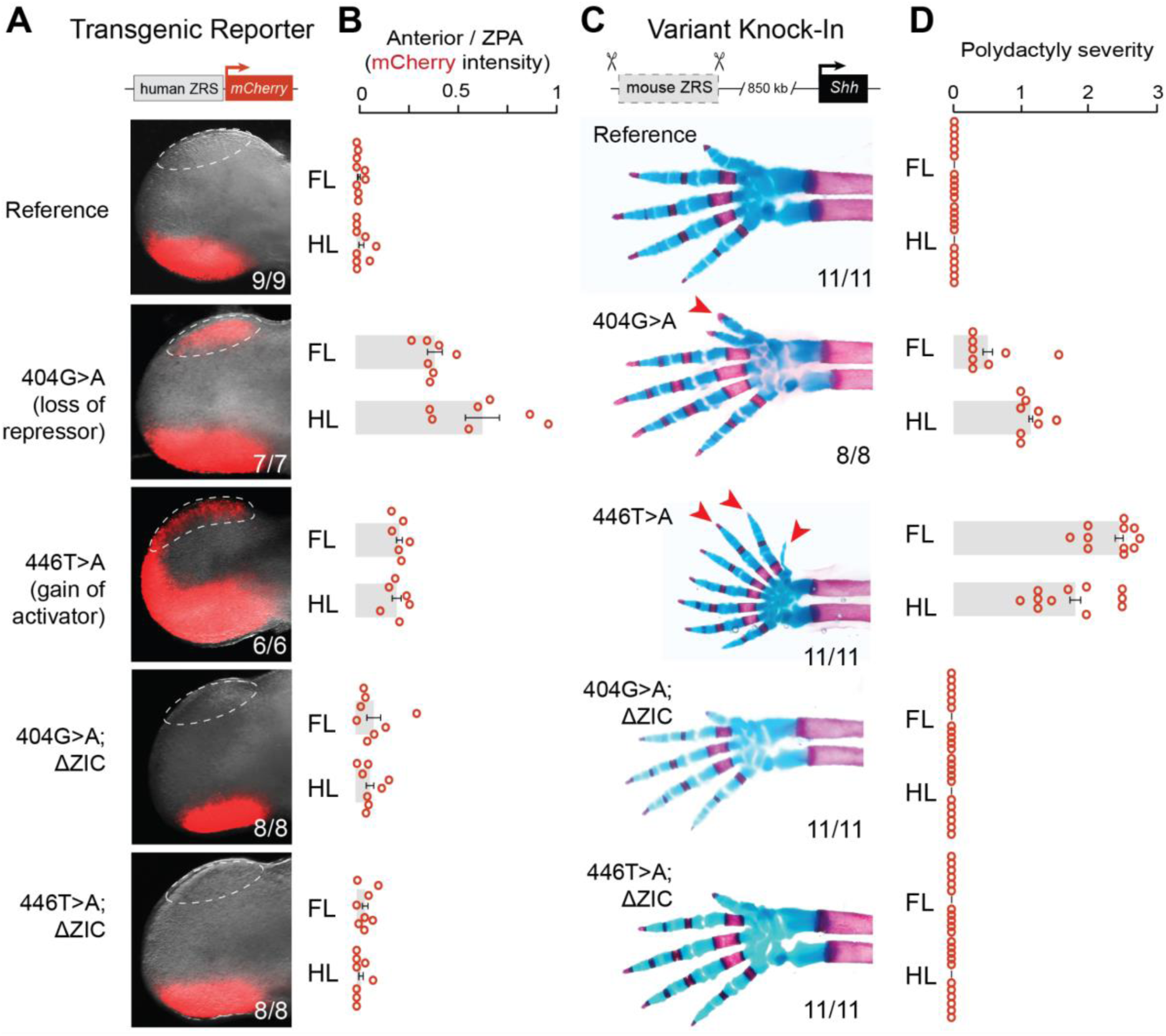
ZIC3 binding site mutagenesis prevents ectopic enhancer activation and rescues limb malformations caused by independent pathogenic variants. (**A**) Fluorescent images of forelimbs from E11.5 transgenic embryos with different human ZRS alleles driving *mCherry*. Numbers indicate independent biological replicates. (**B**) Quantification of relative mCherry intensity in the anterior domain normalized to the ZPA in forelimbs for different hZRS alleles (n = at least 6 independent embryos per genotype). Unpaired student’s t-test with hZRS^ref^: Forelimb (FL): hZRS^404G>A^, *P* < 0.0001; hZRS^446T>A^, *P* < 0.0001; hZRS^404G>AΔZIC^, *P* = ns; hZRS^446T>AΔZIC^, *P =* ns. Hindlimb (HL): hZRS^404G>A^, *P* < 0.0001; hZRS^404G>AΔZIC^, *P* = ns; hZRS^446T>A^, *P* < 0.0001; hZRS^446T>AΔZIC^, *P =* ns. (**C**) Representative skeletal images of forelimbs from knock-in mice in which the native ZRS has been replaced by ZRS alleles carrying human variants and the mutagenized ZIC2/3 motif (n = at least 8 independent mice per genotype). Red, bone; Blue, cartilage. (**D**) Quantification of extra digit number (greater than five) across genotypes. Unpaired student’s t-test with mZRS^Ref^: Forelimb (FL): mZRS^404G>A^, *P* = 0.0015; mZRS^404G>A;ΔZIC2/3^, *P* = ns; mZRS^446T>A^, *P* < 0.0001; mZRS^446T>A;ΔZIC^, *P =* ns. Hindlimb (HL): mZRS^404G>A^, *P* < 0.0001; mZRS^404G>A;ΔZIC^, *P* = ns; mZRS^446T>A^, *P* < 0.0001; mZRS^446T>A;ΔZIC^, *P =* ns.

To assess the extent to which the observed changes in ectopic enhancer activity affect limb morphology *in vivo*, we generated knock-in mice where we simultaneously introduced a human variant and a mutagenized ZIC2/3 motif into the endogenous mZRS enhancer. Mice containing 404G>A or 446T>A variants in the endogenous mZRS have 1-3 extra digits on fore- and hindlimbs as well as tibial hemimelia (shortened or absent tibial bone), mimicking limb malformations observed in patients with these variants (**Figs. 4C, D** and **S7E**) (Wieczorek et al. 2010; Xu et al. 2020; Cho et al. 2013; Zepeda-Olmos et al. 2024). Consistent with the results of transgenic reporter experiments, mice containing 404G>A or 446T>A variants and a mutagenized ZIC2/3 motif in the endogenous mZRS enhancer were phenotypically normal (**Fig. 4C**). All inspected mice had five digits, and their tibia length was not significantly different from wild type mice (**Figs. 4C** and **S7E, F**). Together, these results demonstrate that the ZIC2/3 motif is required for limb malformations induced by independent pathogenic variants. Without ZIC2/3 binding, the ZRS enhancer is essentially resistant to aberrant activation by gain-of-function mutations.

### Spatial poising at developmental enhancers is widespread

We next asked if the observed poised signature is a common feature of mammalian developmental enhancers or, alternatively, if disease-associated loci represent an exception. To address this question we used the VISTA Enhancer Browser, a unique resource of nearly 1,600 human and mouse developmental enhancers with *in vivo* activities experimentally validated in transgenic mice (Pennacchio et al. 2006; Visel et al. 2006; Kosicki et al. 2025). We systematically compared tissue-specific activities of these enhancers with their corresponding tissue-specific chromatin states from ENCODE (Gorkin et al. 2020). Surprisingly, we found that 65% of active developmental enhancers are also accessible in at least one tissue in which they are inactive (**Fig. 5A, B**). For example, 25% of active VISTA enhancers are accessible but inactive in the forebrain at E11.5 (**Fig. 5A**). These spatially poised enhancers are also characterized by H3K4me1 levels comparable to those of active enhancers (*P* = ns; **Fig. 5C**) and are marked by H3K27ac, albeit at lower levels than active enhancers (*P* < 0.0001; **Fig. 5C**). We observed a similar poised enhancer signature characterized by open chromatin, H3K4me1 and low levels of H3K27ac in all five examined tissues of E11.5 embryo (forebrain, midbrain, neural tube, limb and heart) (**Fig. S8A-D**). Spatially poised enhancers were enriched for pioneer TF motifs and displayed an increased overall density of pioneer TF motifs in comparison to enhancers whose accessibility is restricted to active tissues (**Figs. 5D** and **S8E)** (Heinz et al. 2010; Yu and Buck 2019; Boller et al. 2016). Altogether, these data show the presence of a widespread poising signature at developmental enhancers during mammalian organogenesis.

**Figure 5.**
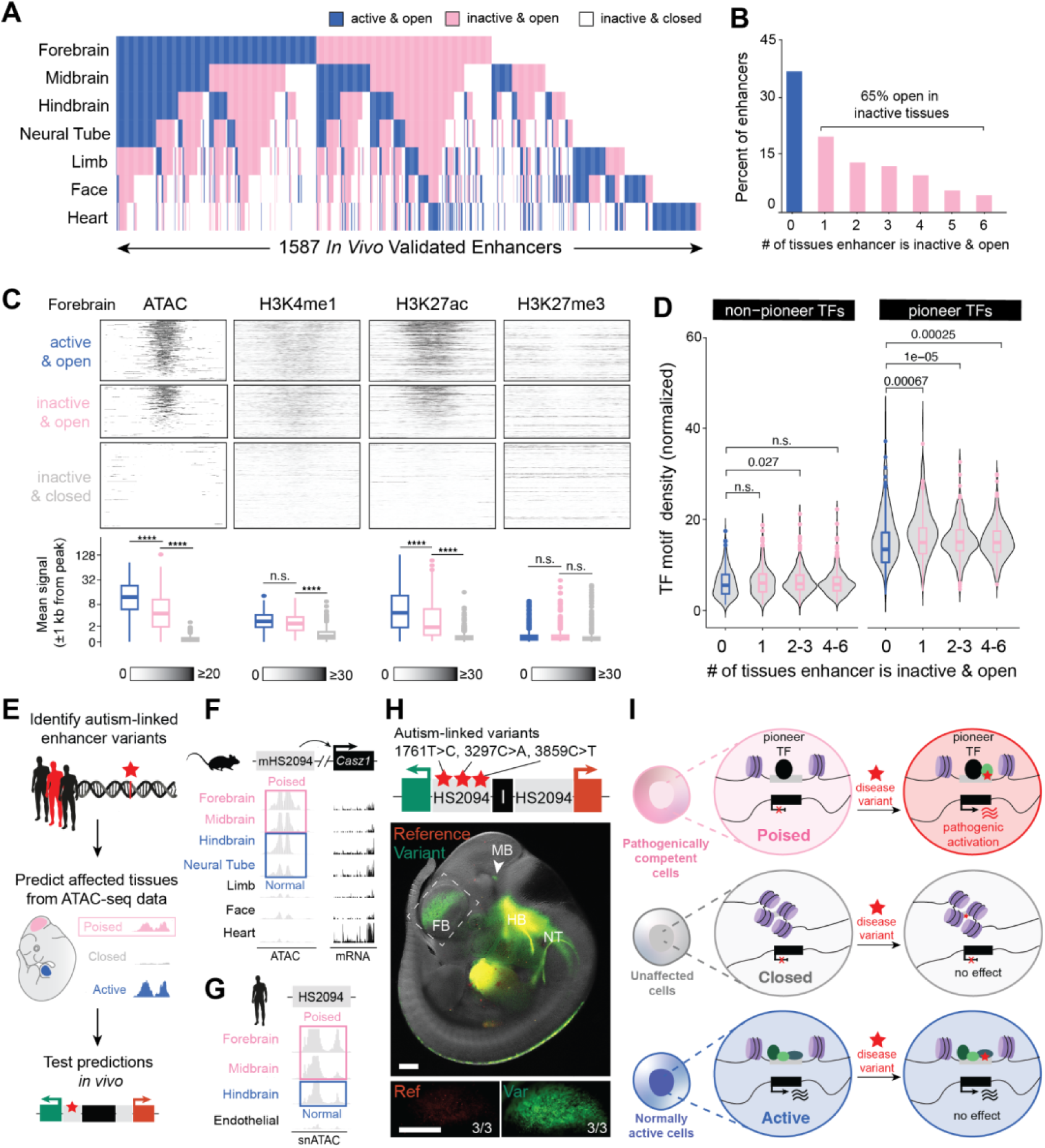
Poising signature predicts the tissue-specific effects of previously uncharacterized non-coding variants associated with autism. (**A**) Heatmap depicting *in vivo* activity (blue) and chromatin accessibility (pink) of nearly 1,600 active VISTA enhancers clustered by their activity across seven embryonic tissues at E11.5. (**B**) VISTA enhancers are classified based on the number of tissues in which they are poised (open and inactive). (**C**) Heatmaps (±2 kb from ATAC peak center) and box plot quantifications of ATAC-seq and ChIP-seq peaks for H3K4me1 and H3K27ac in E11.5 forebrain. Blue, active & open; Pink, inactive & open; Gray, inactive and closed. Unpaired student t-tests. ****, *P* < 0.0001. (**D**) Quantification of the normalized motif density for non-pioneer (left) and pioneer (right) TFs. Unpaired student t-tests. Box plots are median (bold line) ± 25% interquartile range; whiskers extend ±1.5 times the interquartile range. **(E)** Schematic for identifying and testing previously uncharacterized human non-coding variants linked to the neurodevelopmental disorders. **(F)** Chromatin accessibility for mHS2094 and mRNA expression of target gene *Casz1* in bulk across mouse tissues. **(G)** Pseudobulk chromatin accessibility for HS2094 across human fetal brain regions. **(H)** Representative fluorescent image of a transgenic embryo injected with the hs2094 variant allele driving *eGFP* and the reference allele driving *mCherry* (n=3/3). Arrows denote ectopic activity; inset panels, forebrain. Scale bars, 500 μm. **(I)** Summary model depicting how enhancer poising by pioneer factors renders specific cell populations susceptible to aberrant gene activation by non-coding disease variants.

### Poised enhancer signature predicts the effects of autism-linked noncoding variants

Our data suggest that enhancer poising by pioneer factors confers cell-type-specific pathogenic gene activation by noncoding variants at many disease-associated loci. We therefore asked if a poised enhancer signature can predict the tissues and cell types of ectopic activity for previously uncharacterized noncoding variants. To address this question, we compiled a list of all non-coding variants linked to neurodevelopmental disorders identified by GWAS and WGS studies (**Fig. 5E**) (Shin et al. 2024; Wright et al. 2023; Turner et al. 2017; Short et al. 2018). We intersected this list with previously validated in vivo human and mouse enhancers that are active at embryonic day E11.5. To further narrow the list, we focused on enhancers that contained multiple independent, rare human variants and displayed accessible chromatin in tissues outside of their normal domains of activity, reasoning that these regions were the least likely to be random benign mutations (**Figs. 5F and S9**). From this prioritization, we selected ten independent rare variants associated with neurodevelopmental disorders, distributed among four different enhancers. To functionally test these variants, we generated compound variant alleles (containing 2-3 variants per enhancer) for each of the enhancers and placed them upstream of *eGFP* together with the corresponding reference enhancer alleles driving *mCherry* within the same dual-enSERT construct (**Fig. 5E**).

After zygote microinjection, we observed a gain of enhancer activity for HS2094 and HS280 enhancers and no detectable changes for HS1351 and hMM996 enhancers (**Fig. S9A-H**). For example, the HS2094 enhancer of the autism-linked gene *Castor Zinc Finger 1* (*CASZ1*) is normally active in the developing hindbrain and neural tube in mice and contains three rare variants of unknown functional significance (**Fig. 5F**) (Z. Chen et al. 2024; T. Wang et al. 2020; Shin et al. 2024; Zhou et al. 2022). HS2094 is accessible and active in the hindbrain and neural tube, but it is accessible yet inactive in the forebrain and midbrain in mouse and human (**Fig. 5F, G**). Consistent with our prediction, the introduction of three rare, autism-linked 1761C>T, 3297C>A, and 3859C>T variants in the HS2094 enhancer of *CASZ1* resulted in reproducible and strong gain of enhancer activity in the forebrain (3/3 embryos; *P* = 0.0048; **Figs. 5H** and **S9G, H**) with one embryo of the three showing additional ectopic activity in the midbrain (**Figs. 5H** and **S9G, H**). These experiments demonstrate that the poised enhancer signature can be used to inform the *in vivo* effects of candidate non-coding enhancer variants associated with congenital disorders (**Fig. 5I**).

## Discussion

In the present study, we uncovered a cell-type-specific poised enhancer signature that is necessary for ectopic limb-specific *Shh* enhancer activation by pathogenic non-coding variants. We identified a similar cell-type-specific poised signature characterized by open chromatin, H3K4me1, and low H3K27ac at all other human enhancers with known gain-of-function mutations and disease-associated variants. This discovery reveals a property of developmental enhancers that makes some of them susceptible to aberrant activation by non-coding mutations, resulting in disease **(Fig. 5I**).

Canonical poised enhancers—marked by open chromatin, H3K4me1, and sometimes H3K27me3—are primed for future activation, which enables rapid activation of gene expression during lineage specification (Rada-Iglesias et al. 2011; Crispatzu et al. 2021; Bogdanovic et al. 2012; Bonn et al. 2012; Wamstad et al. 2012; Calo and Wysocka 2013; Creyghton et al. 2010) and in response to external cues (Zaret, Lerner, and Iwafuchi-Doi 2016; Beck et al. 2024). The poised signature described in our study is similar, but it occurs along the spatial axis during mammalian organogenesis, as the diversity of tissues and cell types rapidly expands. Such poised signature have been observed during tissue patterning in the early *Drosophila* embryo (Koenecke et al. 2017; Bonn et al. 2012; Junion et al. 2012) and during endodermal differentiation *in vitro* (A. Wang et al. 2015). These spatially poised enhancers could be primed for future activation or they could reflect the developmental history of a cell’s lineage (Koenecke et al. 2017; A. Wang et al. 2015; Junion et al. 2012). The latter possibility seems more likely for the ZRS because *Shh* is never active in anterior limb bud cells during normal development.

Our findings provide a plausible mechanistic explanation for the excess of gain-of-function enhancer variants linked to human disease (**Fig. S1A, B**). While the organism tolerates potential loss-of-function (LOF) enhancer mutations due to substantial intra-enhancer (Crocker et al. 2015; Kvon et al. 2020; Snetkova et al. 2021; Kosicki et al. 2024; Canver et al. 2015) and inter-enhancer (Osterwalder et al. 2018; Kvon et al. 2021; Cannavò et al. 2016) redundancy, gain-of-function enhancer mutations are more likely to affect gene expression because accessible chromatin sensitizes enhancers for ectopic activation.

Our results also have implications for the functional interpretation of the thousands of noncoding variants associated with human disease. It offers a strategy for narrowing the vast biological search space in which a variant may affect gene expression; that is, leveraging the rapidly growing wealth of epigenomic datasets, such as single-cell ATAC-seq (Ziffra et al. 2021; Mannens et al. 2023; Y. E. Li et al. 2023), to identify potentially affected cell types and time points with poised chromatin at a putative non-coding disease locus. Spatially poised chromatin can also be beneficial, as it provides a straightforward mechanism for evolutionary innovation, where a single point mutation can result in a new domain of gene expression (Prud’homme et al. 2006; Gompel et al. 2005; Rebeiz and Tsiantis 2017).

## Acknowledgments

The authors would like to acknowledge the UCI Transgenic Mouse Facility for help with the generation of the enhancer knock-in and transgenic mice, and the Institute for Immunology and Sue & Bill Gross Stem Cell Research Center for their technical assistance with cell sorting. We thank Alexander Stark (IMP), Ivan Marazzi (UCI), Katrina Woolcock (Life Science Editors) and members of the Kvon lab for manuscript editing suggestions. This work was made possible, in part, through access to the Genomics Research and Technology Hub Shared Resource of the Cancer Center Support Grant (P30CA-062203) at the University of California, Irvine and NIH shared instrumentation grants 1S10RR025496-01, 1S10OD010794-01, and 1S10OD021718-01. This work was supported by a National Institutes of Health grant R01HD115268 (to E.Z.K.). E.W.H. was supported by predoctoral fellowships (F30HD110233, T32NS082174, and T32GM008620) from the National Institutes of Health.

## Author contributions

E.W.H. and E.Z.K. conceived the study. E.W.H., C.X.C., T.A.L., and S.H.J. performed transgenic mouse and molecular cloning experiments. E.W.H performed transcriptomic and epigenomic experiments and analyzed the data. E.W.H. and Z.C. performed poised enhancer analysis on ENCODE data. E.W.H., C.X.C, and T.A.L. performed skeletal staining experiments. E.W.H. and E.Z.K. wrote the manuscript with input from all authors.

## Data accessibility

Raw and processed sequencing data are deposited under the following GEO accession numbers: GSE299694 (ATAC-seq), GSE299695 (CUT&Tag), GSE299697 (CUT&RUN), GSE299698 (RNA-seq).

## Methods

### Ethics statement

All animal procedures, including those related to the generation of transgenic and knock-in animals, were conducted in accordance with the guidelines of the National Institutes of Health (NIH) and approved by the Institutional Animal Care and Use Committee at the University of California, Irvine under protocol numbers AUP-20-001 and AUP-23-005.

### Molecular cloning

Enhancer-reporter constructs were generated using previously published reporter plasmids (enSERT mCherry reporter, Addgene #211940 and dual-enSERT reporter, Addgene #211942) (Hollingsworth et al. 2025). All reference enhancer alleles were cloned from human genomic DNA (Promega, G304A) using Q5 High-Fidelity Polymerase (NEB, M0491) or KOD polymerase (Toyobo, #KMM-201). Variant enhancer alleles were synthesized as gBlocks (Integrated DNA Technologies). Reference and variant enhancer alleles were inserted into plasmid backbones digested with either NotI (NEB, R3189) or AgeI (NEB, R3552), respectively, using Gibson-based assembly methods (NEB, E2611) (Gibson et al. 2009). Mouse ZRS knock-in constructs were generated using a previously published targeting vector (Kvon et al. 2020; Bower et al. 2024a). Variant mouse ZRS alleles were synthesized as gBlocks (Integrated DNA Technologies) and inserted into the targeting vector using Gibson assembly (Osterwalder et al. 2022). Restriction enzyme digestion, Sanger sequencing (Retrogen), and whole-plasmid sequencing (Plasmidsaurus) were used to validate sequence fidelity and plasmid identity prior to microinjection. Primers for all cloned sequences are detailed in Table S5.

### Transgenic and knock-in mouse generation

All transgenic and knock-in mice in this study were generated using a CRISPR/Cas9 microinjection protocol, as previously described (Kvon et al. 2020; Osterwalder et al. 2022; Hollingsworth et al. 2025). Briefly, a mix of (i) Cas9 protein (final concentration of 20 ng/μl; IDT Cat. No. 1074181), (ii) sgRNA (50 ng/μl) and (iii) donor plasmid (7 ng/μl) or biotinylated, linearized fragment (1 ng/μl) in injection buffer (10 mM Tris, pH 7.5; 0.1 mM EDTA) was injected into the pronucleus of FVB embryos. All donor plasmids or fragments were column-purified using a PCR purification kit (Qiagen) and eluted into injection buffer before zygote microinjection. Female mice (CD-1 strain) were used as surrogate mothers. Super-ovulated female FVB mice (7–8 weeks old) were mated to FVB stud males, and fertilized embryos were collected from oviducts. The injected zygotes were cultured in M16 with amino acids at 37°C under 5% CO2 for approximately 2 hr. Afterward, zygotes were transferred into the uterus of pseudopregnant CD-1 females. F0 embryos were either collected at E11.5 or brought to gestation. Genotyping for successful integration at the H11 locus (reporter) or the mouse ZRS locus (knock-in) was performed using previously reported primers (Kvon et al. 2016, 2020). PCR- and Sanger-sequencing-based confirmation of ZIC2/3 motif mutagenesis of the mouse ZRS was performed using primers described in **Figure S6** and **Table S5**.

### Mouse maintenance and embryo collection

Mice were housed at a standart housing temperature (19-23°C) in a humidity-controlled (40-60%) facility on a reversed 12-hr dark-light cycle with food and water provided *ad libitum*. Time of gestation was identified by the presence of vaginal sperm plugs, indicating E0.5. Pregnant dams were humanely euthanized, and embryos carefully removed under brightfield stereoscopes in ice-cold PBS (Cytiva, SH30256.01). Yolk sacs or ear punches were collected for genotyping. For embryos injected with batches of enhancer-reporter constructs, PCR genotyping and Sanger sequencing were performed to identify the plasmid identity in each embryo.

### Fluorescent imaging and quantification

Transgenic embryos were imaged in ice-cold PBS in a small petri dish (Greiner Bio-One, #627102) atop a thin layer of 2% gel agarose (Fisher, BP160). Images were taken on a Zeiss V20 stereoscope using a monochromic camera (Axiocam 202, Zeiss), fiber optic light source (Zeiss, CL1500) LED fluorescent laser (X-Cite, Xylis), and fluorescent channels at 488 and 555 nm wavelengths. Single-channel images were merged using Zeiss BioLite software. Quantification of mean fluorescent reporter intensity in the anterior and ZPA was performed by importing the original .czi files into Fiji software (Schindelin et al. 2012). To compare across different fluorophores in embryos injected with dual-enSERT plasmids, mean fluorescent reporter intensities of tissues were quantified, and background fluorescence was subtracted. *Hsp68*-promoter-driven heart fluorescence was used as a negative control for statistical analysis (Hollingsworth et al. 2025).

### Fluorescent-activated cell sorting

Anterior, central and posterior hindlimb bud regions were carefully dissected under a stereoscope in ice-cold PBS, pooled across embryos, and digested with collagenase II (Gibco, #17101015) for 10 min at 700 rpm and 37°C with trituration every 5 min. Digestion was halted by addition of 10% FBS (Thermo Fisher, #A3840201) and dissociated cells were centrifuged at 500g for 5 min. Cells were resuspended in 0.04% BSA (Millipore Sigma, #A1595) and filtered through 70 μM P1000 Flowmi cell filters (SP Bel-Art, #136800040) to remove debris. Cells were sorted into 1.5 mL microcentrifuge tubes using a FACSAria Fusion (BD Biosciences) or FACS Aria II (BD Biosciences) sorter with 70 or 100 μm nozzles at a flow rate no greater than 2.4. Dissociated forebrain tissue was used as a negative control for gating mCherry+ and mCherry-cell populations.

### Flow cytometry-based quantification

After confirming dual-enSERT transgene integration via fluorescent imaging, the anterior domains of E11.5 hindlimb buds were dissected and dissociated into single cells using a previously described protocol (Hollingsworth et al. 2025). The percentage of mCherry+/eGFP+ double-positive cells was then quantified using a FACS Aria II (BD Biosciences) or Novocyte Quanteon (Agilent). Forebrain tissue was used as a negative control for gating background fluorescence. The percentage of double-positive cells shown in **Fig. 2** was calculated as eGFP+/mCherry+ double-positive cells divided by the sum of all fluorescently-positive (mCherry+ only, eGFP+ only, eGFP+/mCherry+) cells. Fisher’s exact tests were used to determine statistical significance of the proportion of double-positive and single-positive cells between alleles.

### Analysis of ENCODE epigenomic and transcriptomic data

For analyses shown in **Figures 1**, **6**, **S1**, **S2**, and **S8**, DNA methylation, ATAC-seq chromatin accessibility, and histone ChIP-seq bigwig files were downloaded from ENCODE (Gorkin et al. 2020). For epigenomic analysis on *in vivo* validated enhancers (**Fig. 6A-C and S8**), signal values of different histone ChIP-seq and ATAC-seq data for embryonic tissues (E11.5) were downloaded from the ENCODE database. Only VISTA enhancers that had at least one ATAC-seq peak were considered in this analysis, resulting in nearly 1600 sequences. These data were further averaged over every 10-bp window. For comparison of the strength of histone marks on enhancers in different categories, signal values were averaged over ± 1 kb from their central coordinates. For motif analysis on enhancers (**Fig.1J**), binding motifs of non-pioneer and pioneer TFs were curated from published papers (Grand et al. 2024; Zhu et al. 2018; Fernandez Garcia et al. 2019) and only those expressed (TPM>0.5) in tissues where enhancers are open were counted, based on tissue-matched RNA-seq data from ENCODE. TF density was calculated by dividing the TF motif number by the length of the ATAC-seq peak. For the ZRS-*Shh* locus analysis (**Fig. S1H**), pseudobulk RNA- and ATAC-seq bigwig files were generated using single-nucleus multi-ome data from E11.5 hindlimb (Bower et al. 2024).

### CUT&Tag

CUT&Tag was performed on fluorescently-purified limb bud cell populations using the CUT&Tag-direct for whole cells with CUTAC protocol, version 4 (Kaya-Okur et al. 2019, 2020; Henikoff et al. 2020). In brief, cells were incubated with concanavalin A-coated magnetic beads (Bangs Laboratories, BP531) for 10 min at room temperature (RT). Bead-bound cells were then incubated for 2 hr at RT with the following primary antibodies: H3K4me1 (CUT&Tag, 1:100; Abcam, ab8895); H3K27ac (CUT&Tag, 1:100; Invitrogen, MA5-23516); ZIC2 (1:100, Abcam, ab150404), or IgG negative control (1:100; Epicypher, #13-0042). After washing, cells were incubated with an anti-rabbit IgG secondary antibody (1:100; Epicypher, #13-0047) for 1 hr at RT. Cells were washed again before binding to the fusion pA/G-Tn5 protein (1:20; Epicypher, #15-1017) for 1 hr at RT. To activate Tn5 and cleave target DNA, samples were incubated in tagmentation buffer and incubated at 37°C for 1 hr. After stopping tagmentation, fragmented DNA was released and proteins degraded by incubating with 1% SDS-ProtK solution at 37°C for 1 hr then 58°C for 1 hr in a thermocycler. Indexing PCR was then performed using previously published i5/i7 primers (Buenrostro et al. 2015) with NEBNext HotStart DNA Polymerase (NEB, ME541L). Indexed DNA was then purified using SPRIselect beads (Beckman Coulter, B23318). Tapestation (Agilent) and Qubit were used to determine DNA fragment sizes and concentrations, respectively. Each sample contained a minimum of 35,000 cells and at least two biological replicates were performed. Samples were paired-end, 150 bp sequenced on an Illumina Nova Seq 6000 or NovaSeq X.

Fastq files from mZRS^WT^, mZRS^404G>A^, and mZRS^446T>A^ mice were aligned to custom mm10 genomes containing an artificial chromosome for the hZRS^404G>A^-mCH or hZRS^446T>A^-mCH plasmid reporter. For mZRS^446T>AΔZIC^ samples, files were aligned to a custom mm10 build with the mutagenized ZIC3 motif and disease variant knocked-in at the mZRS as well as an artificial chromosome for the hZRS^446T>A^-mCH plasmid reporter. To ensure no cross-mapping occurred between the endogenous mZRS on chromosome 5 and the hZRS transgene on chromosome 11, fastq files were also mapped to only mm10 and only the artificial chromosome. The bedtools coverage command was used to confirm no read count differences when mapped separately versus together (Quinlan and Hall 2010). Read-mapping was performed using Bowtie2 (version 2.5.1) with the following parameters: -end-to-end --very-sensitive --no-unal --no-mixed --no-discordant --phred33 -I 10 -X 700. Samtools, version 1.15.1 was used to generate, sort and index bam files, from which bigwig files were generated using deeptools, version 3.5.1. Bigwig files were visualized using WashU Epigenome Browser (D. Li et al. 2022; Robinson et al. 2011). For ZIC2, genome-wide peaks were called from sorted bam files using MACS2 with the following settings: macs2 callpeak -t input_bam -n sample_name -f BAMPE -g mm -q 1e−5 --keep-dup all – nolambda --nomodel --outdir out_dir (Zhang et al. 2008).

### CUT&RUN

CUT&RUN was performed on dissected anterior and posterior hindlimb cells from wild-type E11.5 embryos using Epicypher CUTANA (#14-1048) kits. Experiments were performed according to the manufacturer’s guidelines, except for modifications to DNA isolation. Briefly, 6 x 10^5^ cells were bound to concanavalin A-coated magnetic beads for 10 min before overnight incubation at 4 °C with the following antibodies: ZIC3 (1:20; Abcam, ab222124), H3K4me3 (positive control; 1:100; Epicypher, #13-0060), and IgG (negative control; 1:100; Epicypher, #13-0042). After washing, cells were incubated with fusion protein A/G-MNase for 10 min at RT. Calcium chloride solution was added to activate MNase activity (2 hr, 4 °C) before stopping the cleavage reaction and releasing solubilized DNA fragments (30 min, 37 °C). Phenol-chloroform extraction was then performed to isolate DNA using previously published protocols (Skene and Henikoff 2017). Between 25-50 ng of purified DNA was used for library construction using the Epicypher CUT&RUN Library Prep Kit (#14-1001). DNA was end-repaired, ligated to adapters, and uracils were selectively excised. To minimize adapter dimerization, samples were incubated with Proteinase K at 56 °C overnight before purification with SPRIselect beads the following day. Purified DNA was indexed for 14 cycles with i5/i7 primers and 2X HotStart PCR polymerase followed by SPRIselect bead purification. Library sizes were quantified using Tapestation (Agilent) and paired-end 150-bp sequenced on a NovaSeq 6000 or NovaSeq X (Illumina) for 100M reads. Fastq files were aligned to the mm10 genome using Bowtie2, version 2.5.1 with the following options: -local --very-sensitive-local --no-unal --no-mixed --no-discordant --phred33 -I 10 -X 700 (Langmead and Salzberg 2012). Samtools, version 1.15.1 was used to generate, sort, and index bam files, from which bigwig files were generated using deeptools, version 3.5.1 (H. Li et al. 2009; Ramírez et al. 2016). Bigwig files were visualized using WashU Epigenome Browser (D. Li et al. 2022). Genome-wide peaks were called from sorted bam files using MACS2 with the following settings: macs2 callpeak -t input_bam -n sample_name -f BAMPE -g mm -q 1e−5 -- keep-dup all –nolambda --nomodel --outdir out_dir (Zhang et al. 2008).

### Omni-ATAC-seq

Fluorescently-purified hindlimb or bulk face cells (50-100k per sample) were lysed by incubation with nuclear lysis buffer for 5 min on ice. Nuclei were pelleted by centrifugation at 4°C for 5 min at 600xg, resuspended in ice-cold wash buffer, and quantified using a Countess II (Fisher Scientific). Library construction was performed using the ATAC-seq protocol (Buenrostro et al. 2013, 2015), with transposition buffer based on the Omni-ATAC protocol (Corces et al. 2017). Nuclei were incubated with Illumina Nextera transposase and transposed DNA was cleaned (Zymo Research, #D4004) and preamplified using Nextera i5/i7 primers with five PCR cycles. Additional PCR cycle number was determined by qPCR. After purification with AMPure XP beads (#A63881, Beckman Coulter), DNA libraries were quantified by qPCR with Kapa SYBR Fast Universal for Illumina Genome Analyzer kit. Library size was determined using the Bioanalyzer 2100 DNA High Sensitivity Chip (Agilent). Libraries were paired-end, 150 bp sequenced using Illumina NovaSeq 6000 for 50M reads. The version of NovaSeq control software used was NVCS 1.6.0 with real-time analysis software RTA 3.4.4.

Fastq files were aligned to custom mm10 genomes containing an artificial chromosome for the hZRS^404G>A^-mCH plasmid or hZRS^446T>A^-mCH reporter. For ΔZIC samples, files were aligned to a custom mm10 build with the mutagenized ZIC3 motif and 446T>A disease variant knocked-in at the mZRS as well as an artificial chromosome for the hZRS^446T>A^-mCH plasmid reporter. To ensure no cross-mapping occurred between the endogenous mZRS on chromosome 5 and the hZRS transgene on chromosome 11, fastq files were also mapped to only mm10 and only the artificial chromosome. The bedtools coverage command was used to confirm no read count differences when mapped separately versus together (Quinlan and Hall 2010). Paired-end reads were first trimmed using Fastp, version 0.23.2 and then mapped using Bowtie2, version 2.5.1 with the following parameters: --local --very-sensitive --no-mixed --no-discordant -I 25 -X 700. Bam files were sorted and indexed using samtools, version 1.15.1, and converted to bigwig files for genome visualization using deeptools, version 3.5.1 (H. Li et al. 2009; Ramírez et al. 2016; Martin 2011).

### RNA-seq

Fluorescently-purified cells were snap-frozen in liquid nitrogen and stored at -80°C. RNA was isolated using the RNeasy extraction kit (QIAGEN), following manufacturer’s guidelines. cDNA libraries were constructed and sequencing performed by Novogene for an average of ∼25M reads per sample. All RNA-seq experiments were performed in biological replicates (at least 2 litters of embryos). Paired-end reads were aligned to the mm10 reference genome using STAR, version 2.7.9a with default parameters (Dobin et al. 2013). Expression values were counted using HTSeq and differential expression analysis performed using DESeq2, version 3.16 (Love, Huber, and Anders 2014; Anders, Pyl, and Huber 2015). Genes were considered differentially expressed if Benjamini-Hochberg-adjusted *P* value < 0.05 and log_2_ fold-change > ± 2. Transcription factors were filtered using AnimalTFDB (W.-K. Shen et al. 2023).

### Motif enrichment analysis

*In vivo* validated human enhancers were dichotomized into being inactive & open (in at least one tissue) or those whose accessibility pattern perfectly matched tissue activity. Sequence coordinates were extracted and motif enrichment analysis was performed between the groups using findMotifsGenome.pl from HOMER with standard parameters except for -size given (Heinz et al. 2010). TF motifs were filtered to those expressed in any E11.5 mouse tissue using ENCODE bulk RNA-seq datasets.

### Motif mutagenesis in transgenic and knock-in mice

To mutagenize the ZIC2/3 motif within the hZRS and mZRS, the cognate motif sequence was randomized. The ZIC2/3 motif was mutated from CCTGCTG to AATGAGA. FIMO and HOMER were both performed on mutagenized sequences to validate that no other TF motifs were recognized (Heinz et al. 2010; Grant, Bailey, and Noble 2011). Custom DNA sequences with the mutated motif spanning the entire ZRS length were synthesized as gBlocks (IDT, San Diego, CA) and assembled into plasmids using Gibson-based assembly techniques.

### Skeletal staining

Skeletal staining was performed on P0 mouse pups as described previously using Alcian blue (Sigma, A-3157) and Alizarin red (Sigma, A-5533) to stain cartilage and bone, respectively (Ovchinnikov 2009; Kvon et al. 2016, 2020). Stained fore- and hindlimbs were dissected in 80% glycerol and imaged using a ZEISS Stemi 508 microscope and Axiocam 208 digital camera. Images were processed using Zen BioLite software (ZEISS). Polydactyly severity was scored blind to genotype. Severity was considered zero if all five digits were present, 0.5 for each digit that was bifurcated (only cartilage), and 1 for each digit that was ossified. The severity score for fore- and hindlimbs was averaged per mouse.

### Statistical analysis and reproducibility

Statistical analyses were performed using R version 4.3.1 and Microsoft Excel version 16.79.1. No statistical methods were used to predetermine sample size nor was randomization performed. The investigators were blinded to genotype for all imaging and quantification analyses. No data were excluded from the analyses. P-values (or FDR when applicable) less than 0.05 were considered significant. For all sequencing-based experiments, at least two biological replicates were performed and analyzed without blinding, per field standard. Raw sequencing data were analysed on the UCI high-performance computing cluster.

## Supplementary Figures

**Supplementary Figure 1:**
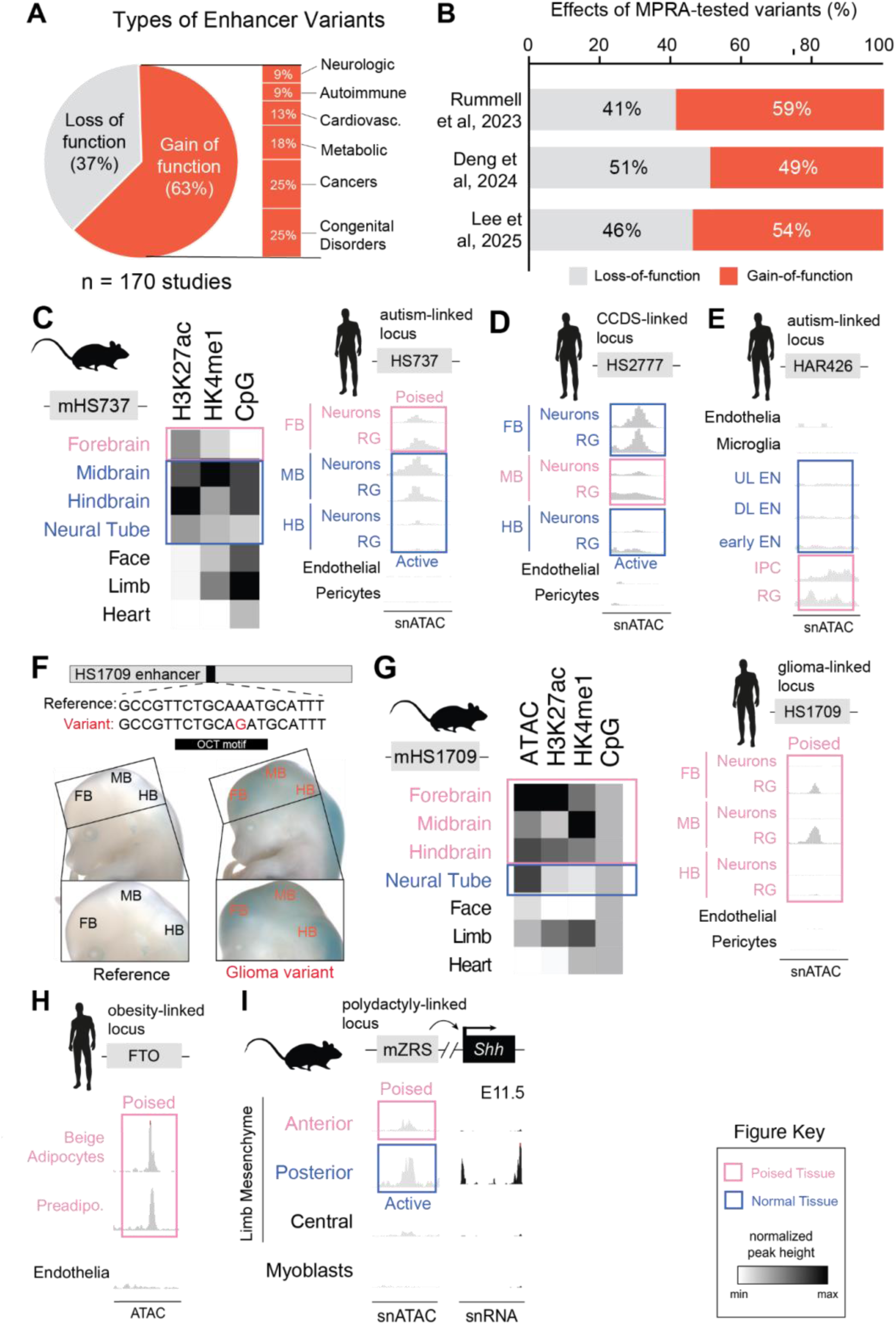
Tissue-specific epigenomic states of disease-linked human enhancers. **(A)** Pie chart depicting distribution of disease-associated enhancer variants reported in the literature based on their type (n=170 independent studies; **Table S1**). On right, breakdown for gain-of-function SNVs. **(B)** Bar plot showing the effects of human disease-associated enhancer SNPs from published MPRA studies. **(C)** Heatmap depicts normalized peak heights for ATAC-seq, ChIP-seq for H3K27ac, H3K4me1, and CpG methylation (left to right) for mouse orthologs of autism-associated HS737 enhancer at E11.5. On the right are pseudobulk cell-type-specific accessibility tracks in the developing human fetal brain from 6 to 13 post-conception weeks (Mannens et al. 2023). FB, Forebrain; HB, Hindbrain; MB, Midbrain; RG, Radial Glia. **(D)** Pseudobulk chromatin accessibility across developing human brain cell types at the HS2777 locus linked to congenital cranial dysinnervation syndrome (CCDS) (Mannens et al. 2023). **(E)** Chromatin accessibility for autism-linked HAR426 in cell types of the developing human neocortex. DL EN, Deep layer excitatory neuron; IPC, Intermediate progenitor cell; UL EN, Upper layer excitatory neuron. Human snATAC-seq data derived from (Trevino et al. 2021). **(F)** Transgenic E14.5 whole embryos and brains in which a human reference HS1709 enhancer allele (left) and a glioma-linked HS1709 variant allele (right) drive LacZ (Yanchus et al. 2022). FB, Forebrain; MB, Midbrain; HB, Hindbrain. **(G)** Same as in (C) but for mHS1709 and HS1709. **(H)** Bulk chromatin accessibility in endothelial cells, preadipocytes and differentiated adipocytes for the obesity-linked FTO locus. Data derived from (Wachowski et al. 2024). **(I)** Genomic tracks of single-nucleus chromatin accessibility and gene expression for mZRS (mouse homolog) and its target gene *Shh*, respectively, from wild type E11.5 hindlimb (Bower et al. 2024).

**Supplementary Figure 2:**
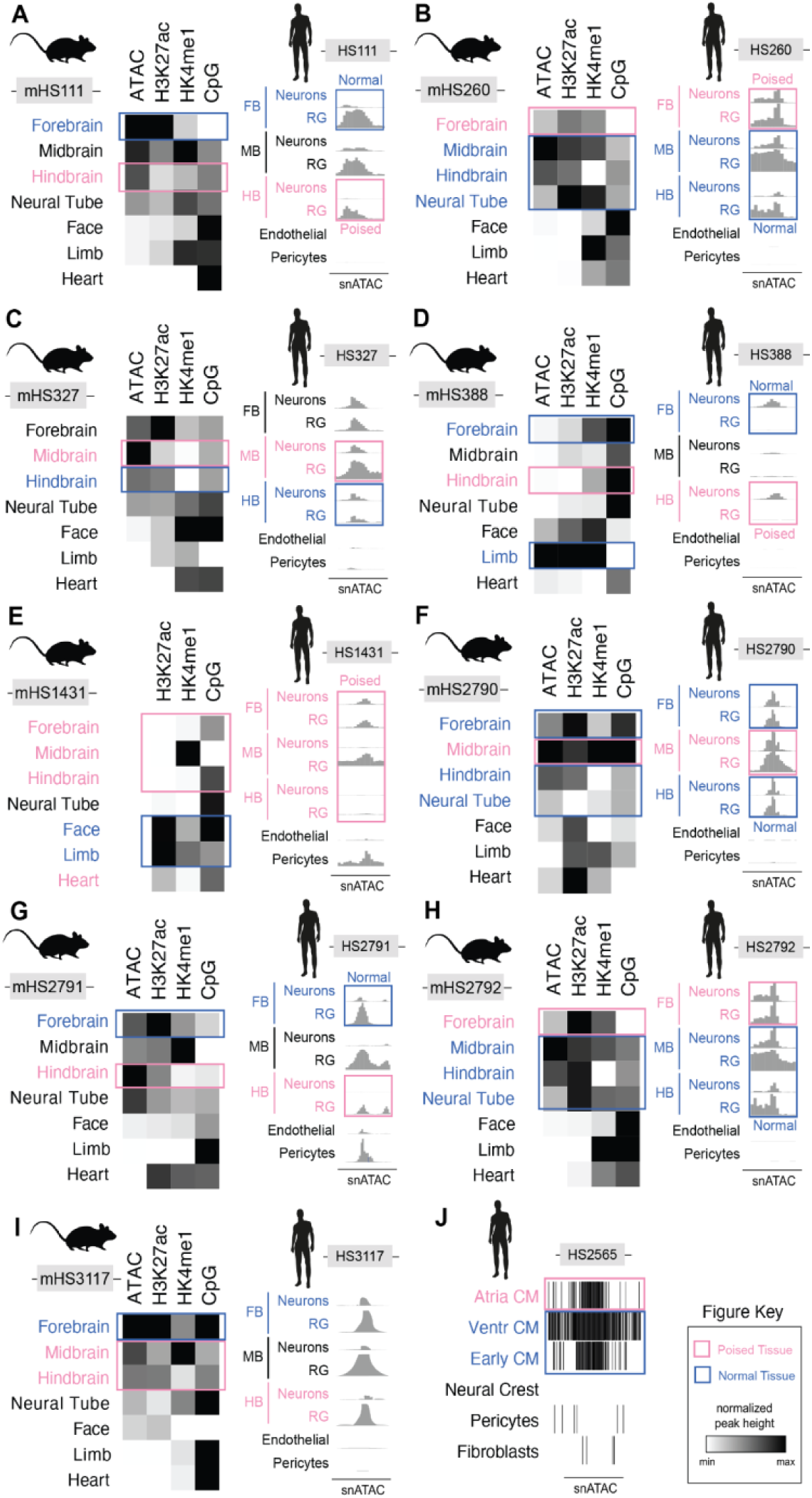
Epigenomic states of human enhancers containing known artificial gain-of-function mutations in mouse and human tissues and cell types. **(A-I)** Heatmaps depict normalized peak intensity for ATAC-seq, histone modifications H3K27ac and H3K4me1, as well as CpG methylation signal in mice for mHS111 (A), mHS260 (B), mHS327 (C), mHS388 (D), mHS1431 (E), mHS2790 (F), mHS2791 (G), and mHS2792 (H), and mHS3117 (I). Shown on right are pseudobulk cell-type-specific accessibility tracks across human fetal brain cell types from publicly-available single-nucleus ATAC-seq data (Mannens et al. 2023). FB, Forebrain; HB, Hindbrain; MB, Midbrain; RG, Radial Glia. **(J)** Pseudobulk chromatin accessibility of human fetal cardiac cell types. Data derived from human snATAC-seq (Ameen et al. 2022). CM, Cardiomyocytes. Blue, tissues in which a reference enhancer allele is active; Pink, tissues in which ectopic enhancer activity is observed.

**Supplementary Figure 3:**
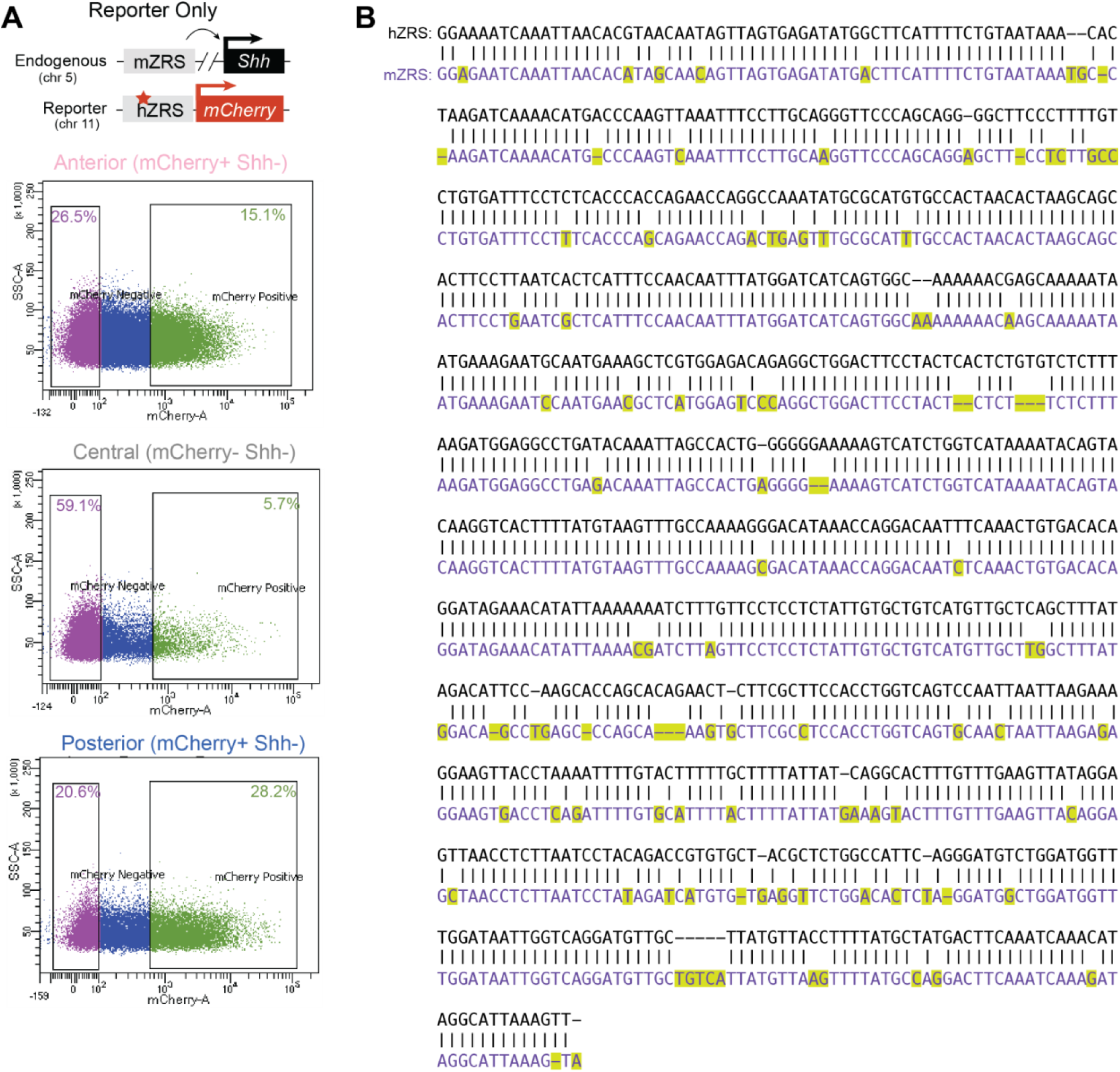
hZRS^404G>A^-reporter cell sorting and mouse-human ZRS alignment. **(A)** Representative fluorescent-associated cell sorting plots across distinct limb bud cell populations in mZRS^WT^; hZRS^404G>A^-*mCH* mouse embryos. **(B)** Sequence alignment of mouse and human ZRS.

**Supplementary Figure 4:**
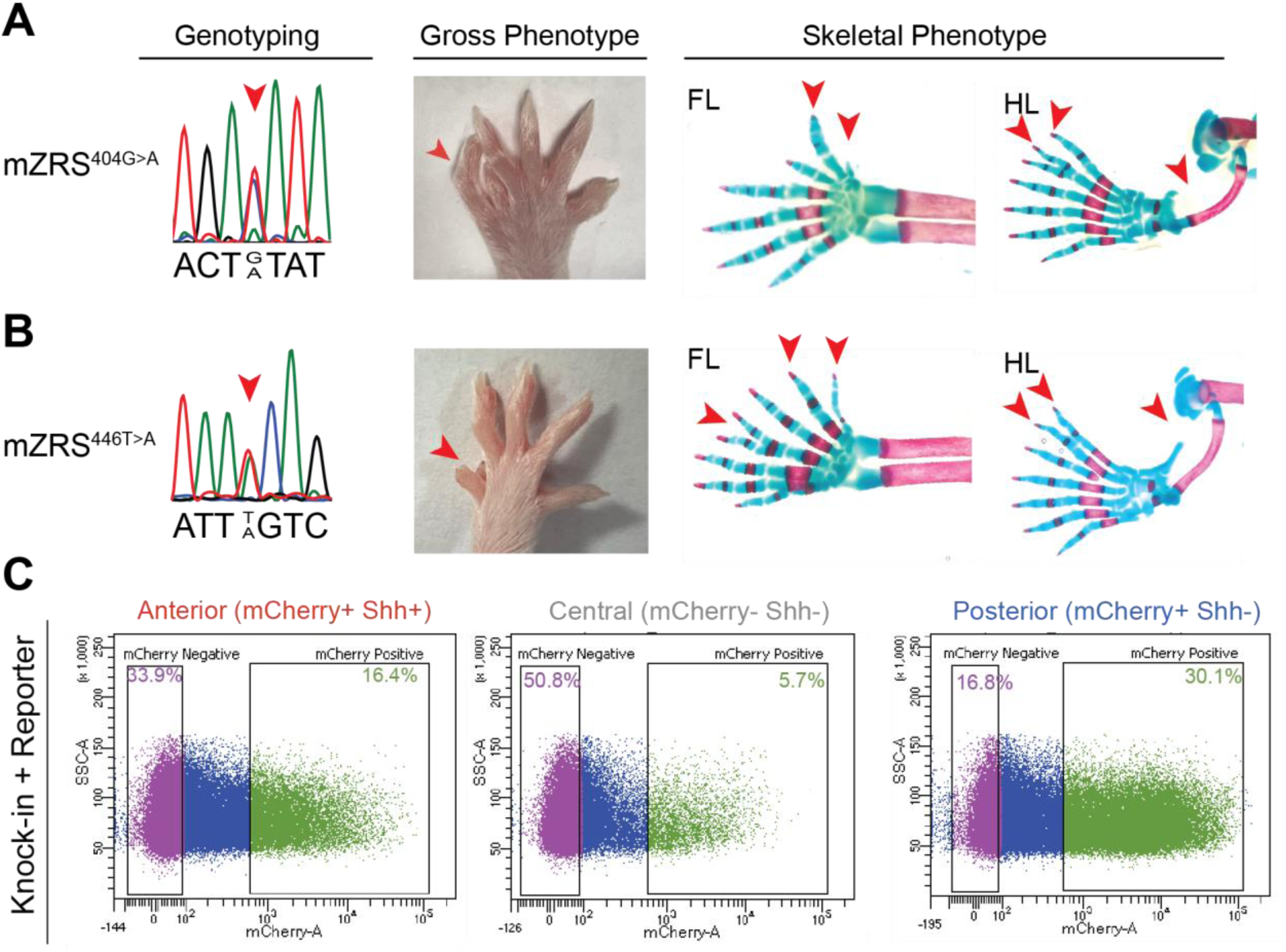
Limb phenotypes of variant knock-in mice. **(A, B)** Sanger sequencing traces, gross limb morphology and skeletal phenotypes of knock-in mice containing 404G>A (A) and 446T>A (B) variants. FL, forelimb; HL, hindlimb. Primers used for genotyping and sequencing are derived from (Kvon et al. 2016, 2020). Note that mice in gross phenotype are F0 heterozygous and those shown in skeletal phenotype are F2 homozygotes. **(C)** Representative FACS plots of limb bud cell populations from 404G>A variant knock-in mouse embryos with a 404G>A variant hZRS reporter transgene driving mCherry (mZRS^404G>A^; hZRS^404G>A^-mCH).

**Supplementary Figure 5:**
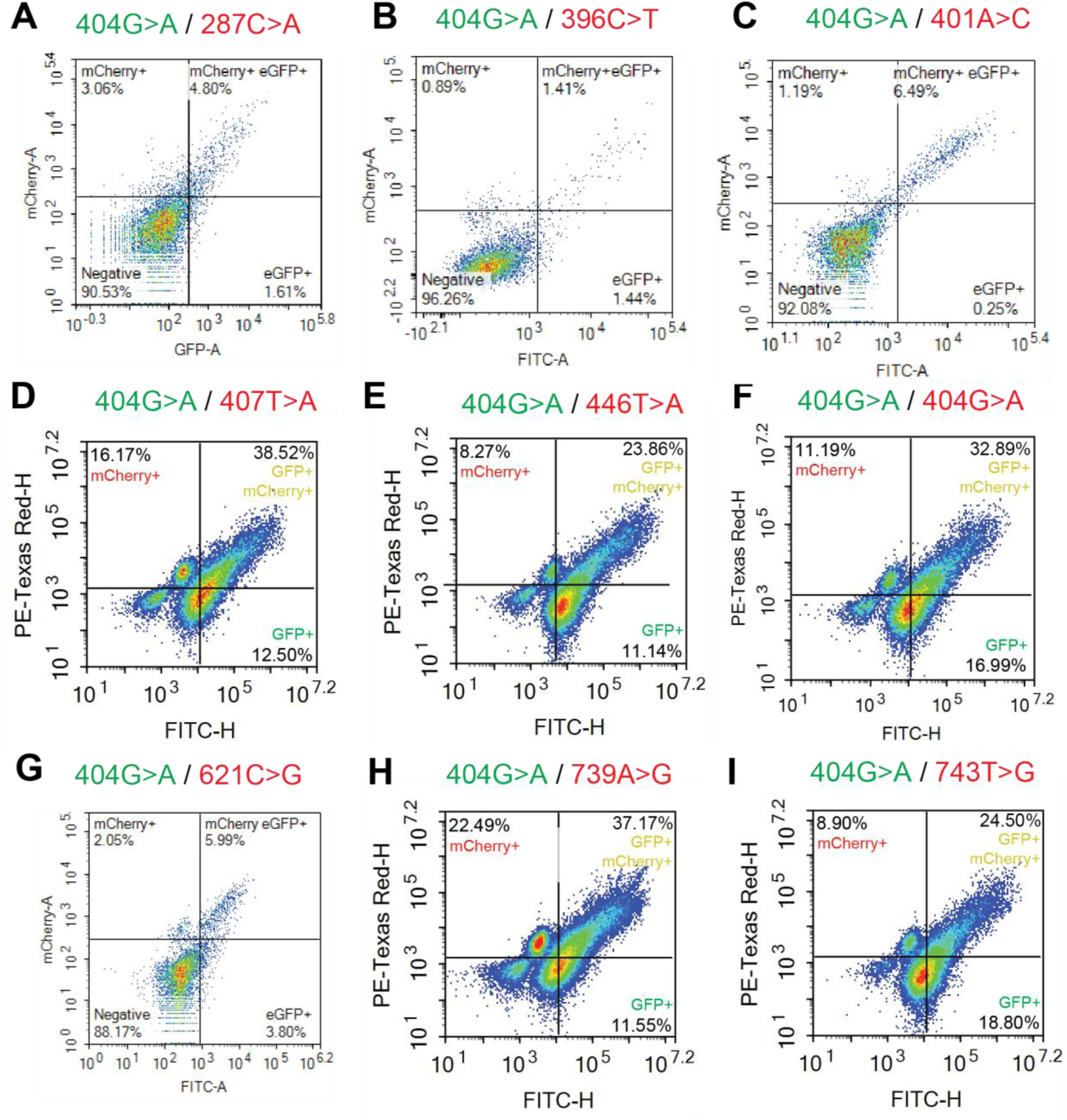
Comparative flow cytometry-based analysis in dual-enSERT transgenic mice. **(A-I)** Flow cytometry plots of anterior portion of hindlimbs from transgenic E11.5 embryos injected with dual-enSERT constructs with hZRS^404G>A^ driving eGFP, and 287C>A (A), 396C>T (B), 401A>C (C), 404G>A (D), 407T>A (E), 446T>A (F), 621C>G (G), 739A>G (H), 743T>G (I) driving mCherry. Note that samples from A-C and G were quantified on a FACS Aria II sorter while samples belonging to D-F and H-I were sorted on a Novocyte Quanteon. Fisher’s exact test against 404G>A/404G>A: 287C>A, *P* = ns; 396C>T, *P* = ns; 401A>C, *P* = ns; 407T>A, *P* = ns; 446T>A, *P* = ns; 621C>G, *P* = ns; 739A>G, *P* = ns; 743T>G, *P* = ns.

**Supplementary Figure 6:**
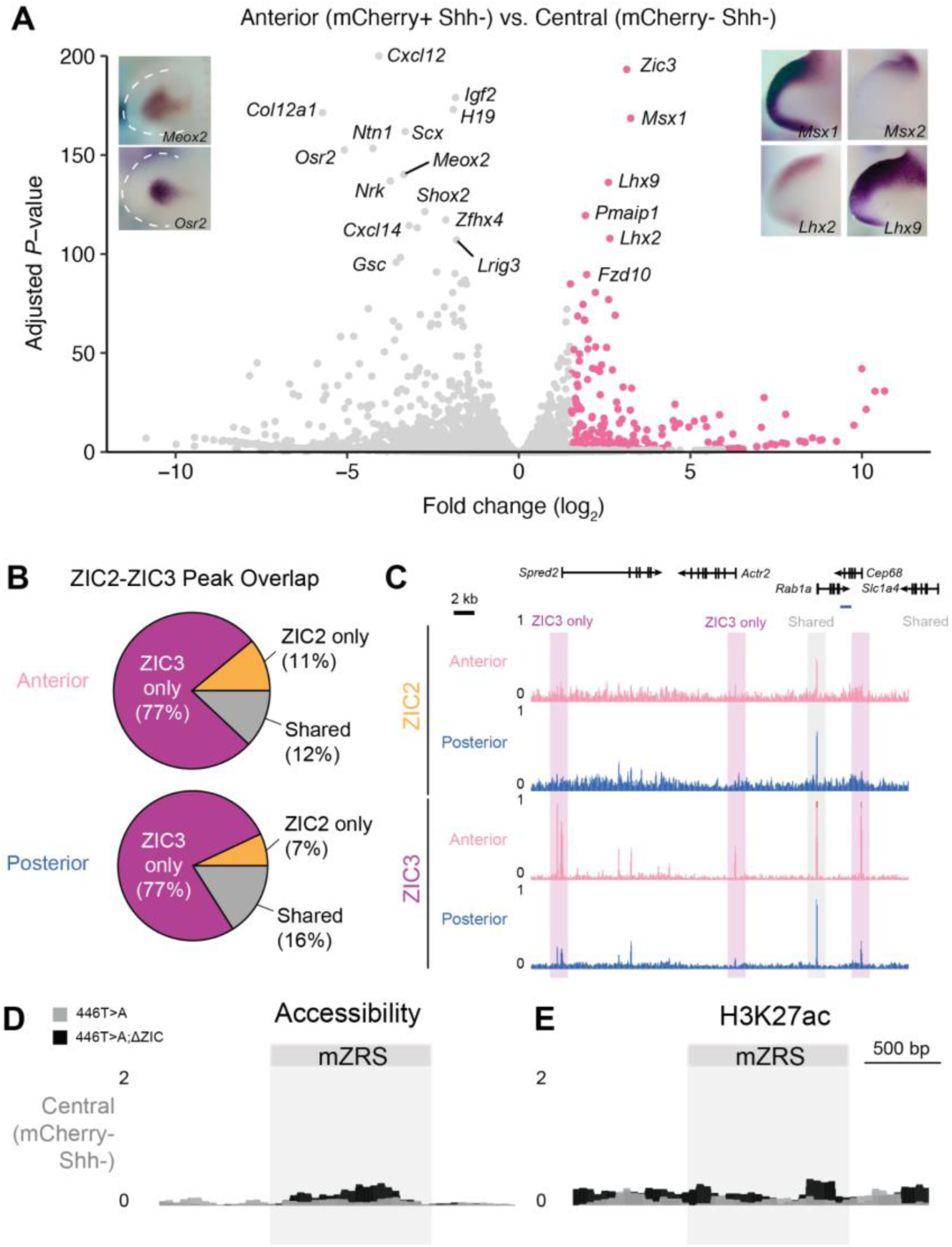
Genome-wide ZIC2/3 binding analysis in different population of limb bud cells. **(A)** Volcano plot for all genes with example *in situ* hybridization images of topmost anterior- (right) and central (right) differentially expressed genes. Images have been reproduced with permission from Embrys database (http://embrys.jp). Pink, significantly upregulated genes in anterior; Grey, insignificant genes. **(B)** Pie charts depict percent overlap of ZIC2 and ZIC3 peaks across anterior and posterior cell populations. **(C)** Genomic tracks for representative examples of ZIC3 only (magenta) and shared ZIC2-ZIC3 peaks (grey). **(D, E)** Genomic tracks for chromatin accessibility (D, ATAC-seq) and H3K27ac (E, CUT&Tag) signal at the mZRS locus in FAC-sorted mCherry-central limb cells (negative control).

**Supplementary Figure 7:**
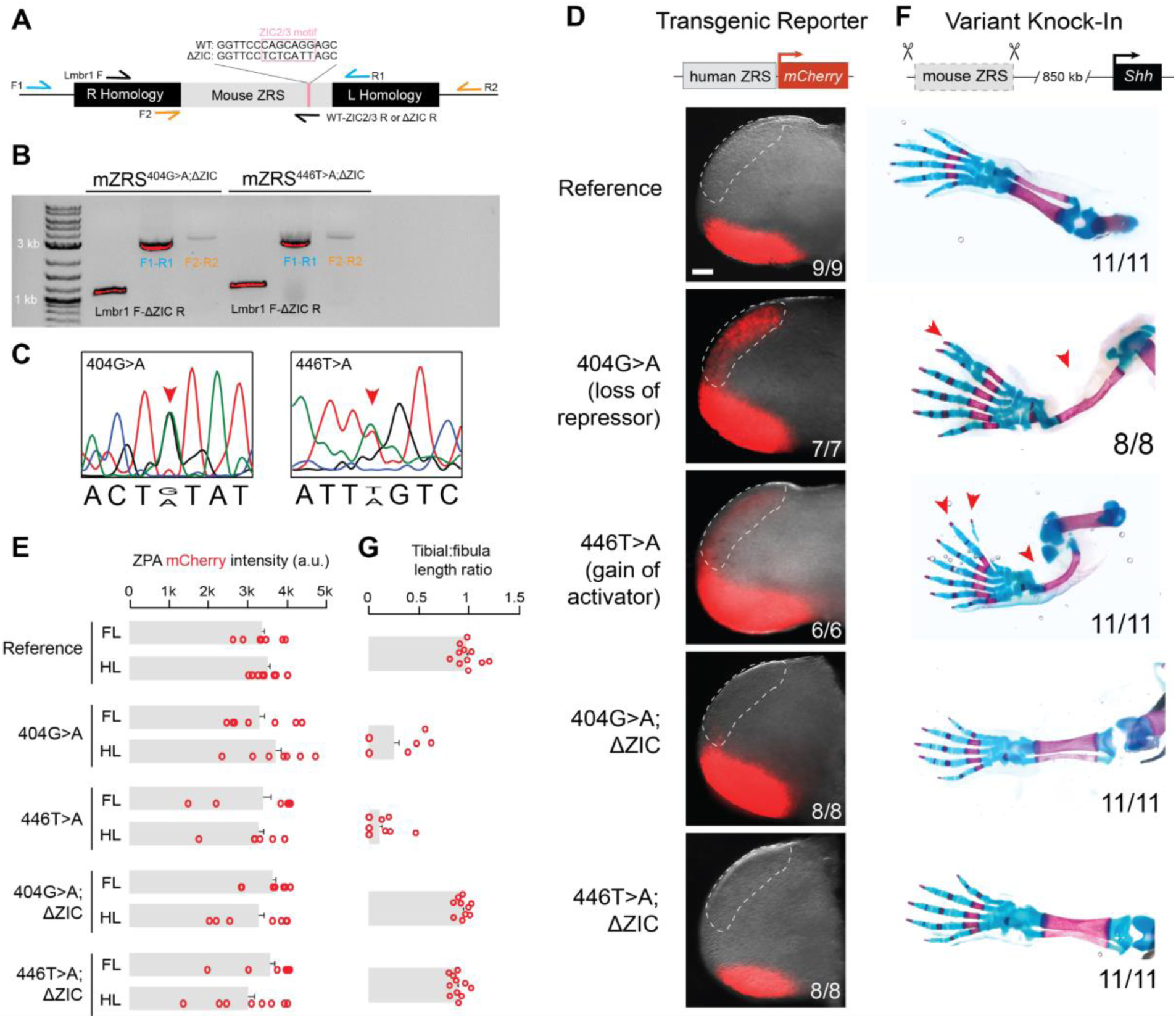
Effect of ZIC2/3 motif mutagenesis on ectopic enhancer activity and limb phenotypes. **(A)** Genotyping schematic for validation of variant mZRS knock-in with a mutagenized ZIC2/3 motif. Forward and reverse primers shown as arrows. ZIC2/3 motif and ΔZIC primers highlighted in pink. Homology arm primer pairs shown in blue and orange. **(B)** PCR gel electrophoresis confirming knock-in of endogenous mZRS with mZRS^404G>AΔZIC^ and mZRS^446T>AΔZIC^. **(C)** Sanger sequencing traces of F0 mice showing heterozygous knock-in of 404G>A and 446T>A variants at the mZRS. **(D)** Sample fluorescent images of hindlimbs from E11.5 transgenic embryos (n/n) with different human ZRS alleles driving mCherry. Quantifications of reporter intensity are included in Figure 4. **(E)** Plot depicting quantification of average mCherry intensity in posterior ZPA of forelimb (FL) and hindlimbs (HL) across genotypes. Unpaired student’s t-test with hZRS^Ref^: All comparisons, *P* = ns. **(F)** Representative skeletal images of hindlimbs across different knock-in mouse lines at the mZRS. Quantifications of digit numbers can be found in Figure 4. **(G)** Plot quantifying ratio of ossified bone lengths in hindlimbs (tibia:fibula) across mouse ZRS knock-in genotypes. Unpaired student’s t-test with mZRS^Ref^: mZRS^404G>A^, *P* < 0.0001; mZRS^446T>A^, *P* < 0.0001; mZRS^404G>A;ΔZIC2/3^, *P* = NS; mZRS^446T>A;ΔZIC2/3^, *P* = NS.

**Supplementary Figure 8:**
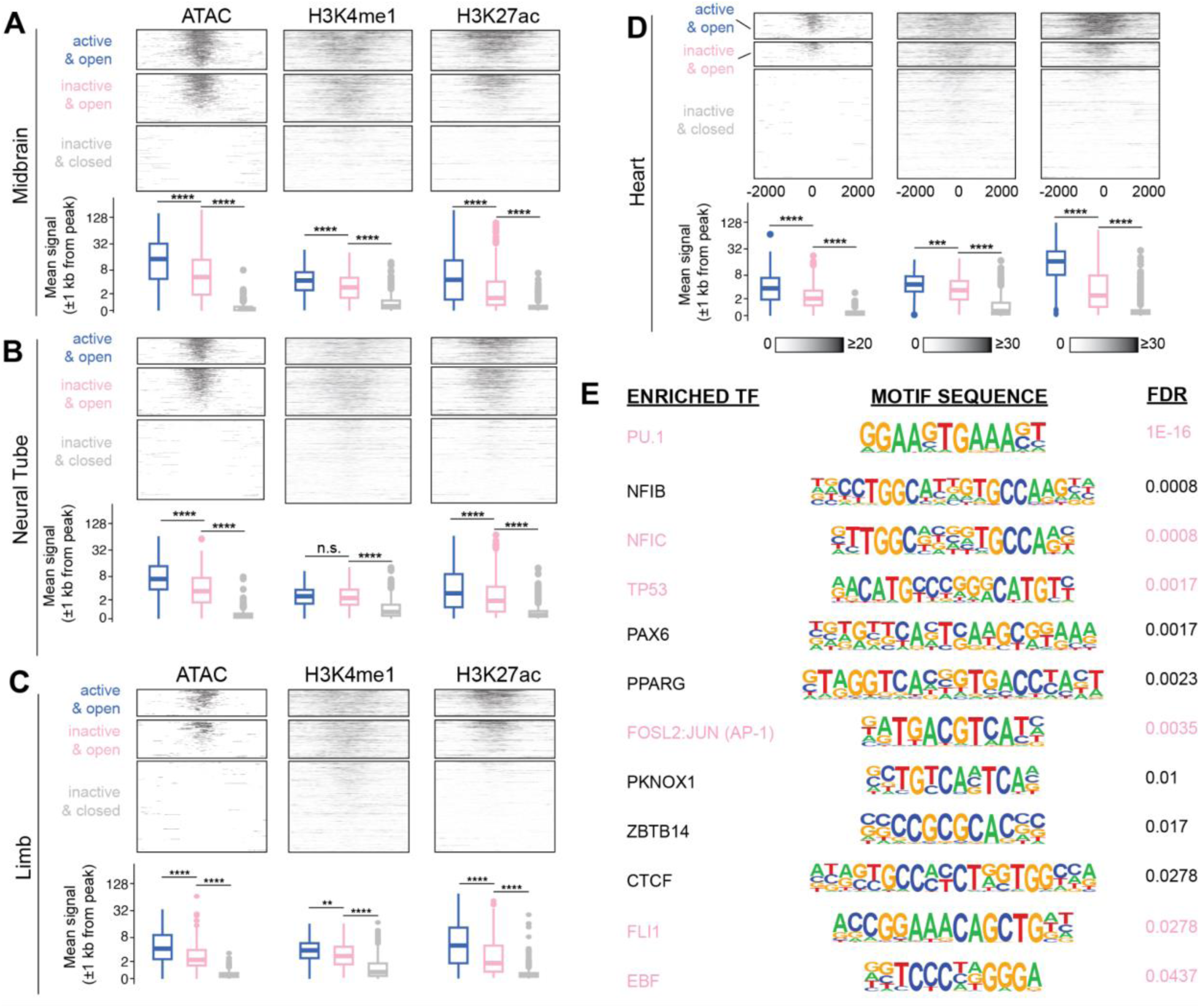
Epigenomic states of VISTA enhancers categorized by tissue-specific activity and accessibility. (**A-D**) Heatmaps of ATAC-seq, H3K4me1, and H3K27ac centered on ATAC-seq peaks, for midbrain (A), neural tube (B), limb (C), and heart (D) at E11.5. Log_2_ box plots show quantification of peak signal across each of the groups. Unpaired student t-tests. Box plots are median (bold line) ± 25% interquartile range. Whiskers are smallest or largest values within 1.5 times of 25th or 75th quartiles. *P* < 0.0001, ****; P < 0.001, ***; P < 0.05, *. Blue, active & open; Pink, inactive & open; Grey; inactive & closed. **(E)** Motif enrichment analysis between poised enhancers and enhancers with active-tissue-restricted accessibility. Pioneer factors are colored in pink. Fisher’s-exact test: Pioneer vs. Non-pioneer motifs, *P* < 0.00001.

**Supplementary Figure 9:**
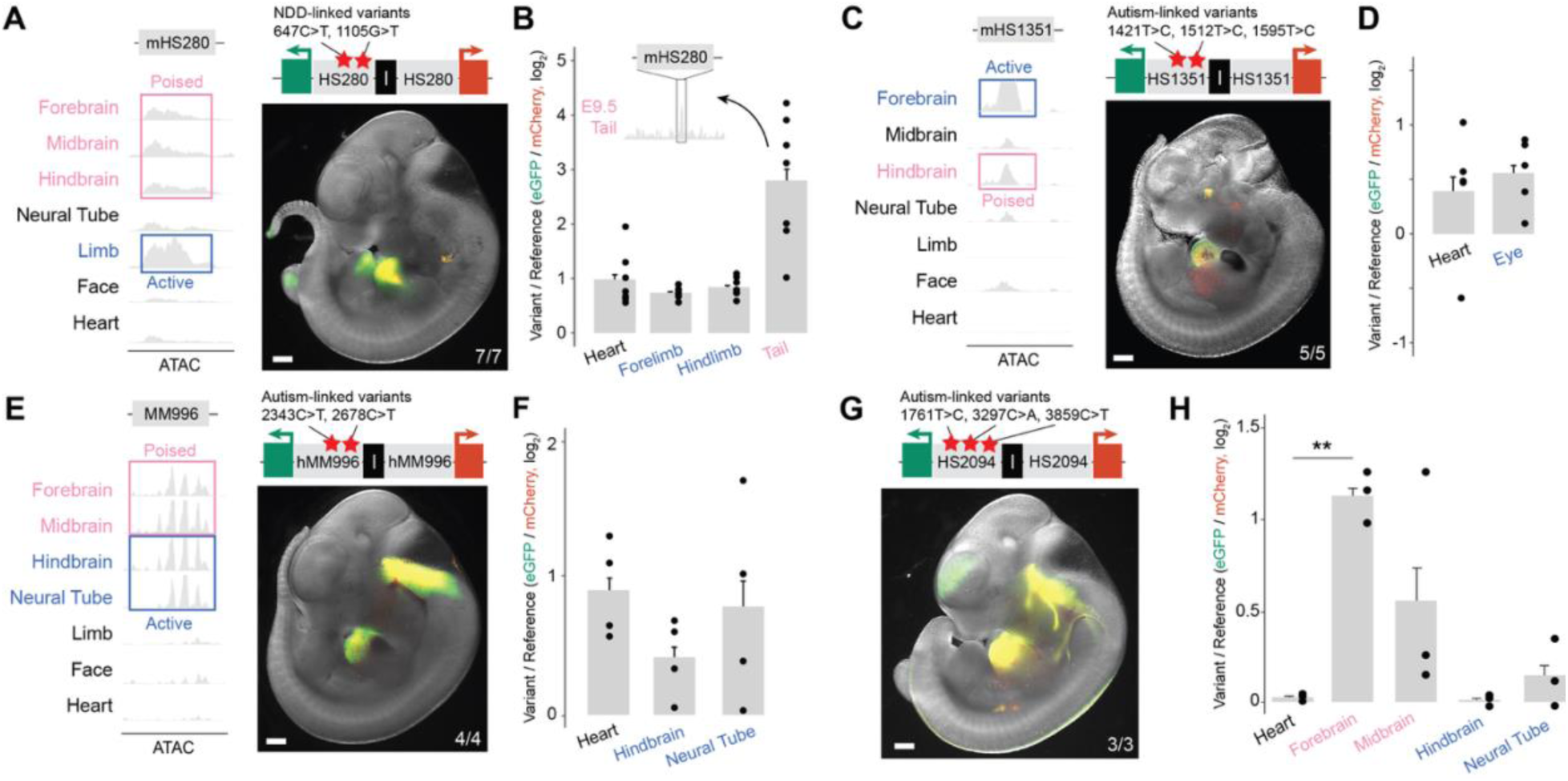
***In vivo* characterisation of enhancer variants associated with autism. (A)** Genomic tracks of bulk-tissue ATAC-seq for mouse ortholog of HS280 and representative images of transgenic E11.5 embryos in which compound variant HS280 allele drives *eGFP* and a reference HS280 allele drives *mCherry*. **(B)** Quantification of fold-change difference (log_2_) between variant and reference enhancer alleles of HS280. Also shown is bulk ATAC-seq data from mouse tailbud at E9.5 for the mHS280 locus (Orchard et al. 2019) where HS280 becomes ectopically active by disease variants. Paired student’s t-test using Heart as negative control: Forelimb, *P* = n.s.; Hindlimb, *P* = n.s.; Tail, *P* = 0.0027 (n = 7 independent embryos). **(C, D)** Same as in (A, B), but for HS1351. Paired student’s t-test using Heart as negative control: Eye, *P* = n.s. (n = 5 independent embryos). **(E, F)** Same as in (A, B), but for hMM996. Paired student’s t-test using Heart as negative control: Hindbrain, *P* = n.s.; Neural Tube, *P* = n.s. (n = 4 independent embryos). **(G, H)** Same as in (A, B), but for HS2094. Paired student’s t-test using Heart as negative control: Forebrain, *P* = 0.0048; Midbrain, *P* = ns; Hindbrain, *P* = ns (n = 3 independent embryos).

## Supplementary Tables

**Table S1.** Classification of human enhancer variants reported in the literature based on their effect on enhancer activity. Only enhancer variants experimentally tested by reporter-based assays, such as lacZ or luciferase, or functional variant knock-ins were considered.

**Table S2.** Human enhancers containing variants and artificial mutations that cause ectopic activity.

| Enhancer | Coordinates (hg38) | Variant(s) | Disease association | Putative target gene(s) | Normal activity | Ectopic activity | CTCF site? | Reference |
| --- | --- | --- | --- | --- | --- | --- | --- | --- |
| HS2777 | chr17: 48002673-48004553 | chr17:48003393G>A, chr17:48003557C>G, chr17:48003752A>C, chr17:48003826C>T | Congenital cranial dysinnervation syndrome (CCDS) | *CDK5RAP3* | Cranial nerves, Forebrain, Midbrain, Hindbrain, Neural Tube | Expansion of Midbrain, Hindbrain | No | Lee et al, Nat Comm 2024 |
| HS737 | chr10: 128568604-128569741 | chr10:128568697A>G, chr10:128569417T>C, chr10: 128569437G>A | Autism | *EBF3* | Neural Tube, Hindbrain, Midbrain | Forebrain | No | Turner et al, Cell 2017; Padhi et al, Hum Genom 2021 |
| HS1709 | chr8: 129631847-129635071 | chr8:129633446A>G | Low-grade glioma | *MYC* | Neural Tube | Forebrain, Midbrain, Hindbrain | No | Yanchus et al, Science 2022 |
| HAR426 | chr7: 101606017-101606400 | chr7:101606359G>A | Autism | *CUX1* | Layer 2/3 cortical neurons | All cortical layer cells | No | Doan et al, Cell 2016 |
| FTO | chr16: 53703963-54121941 | chr16:53767042T>C, chr16: 53779455G>T, chr16:53786615A>T | Obesity | *IRX3, IRX5* |  | Preadipocytes, Beige adipocytes | No | Claussnitzer et al, NEJM 2015 |
| chr21q22 | chr21: 39092644-39096644 | chr21:39094644G>A | Autoimmune disorders | *ETS2* | Monocytes | Macrophages | No | Stankey et al, Nature 2024 |
| HS111 | chr7: 42152129-42154039 | 5% mutagenesis | - | *GLI3, PSMA2* | Forebrain, Neural tube | Hindbrain | No | Snetkova et al, Nat Genet 2021 |
| HS327 | chr1: 88461113-88462825 | 5% mutagenesis | - | *LMO4* | Forebrain, Hindbrain, Neural Tube | Midbrain | No | Snetkova et al, Nat Genet 2021 |
| HS388 | chr2: 7634262-7634939 | 2% mutagenesis | - | *ID2* | Forebrain, Limb | Hindbrain | No | Snetkova et al, Nat Genet 2021 |
| HS260 | chr4: 104424418-104425738 | 5% mutagenesis | - | *GSTCD, INTS12, PPA2, TET2, CXCC4* | Hindbrain, Midbrain, Neural Tube | Forebrain | No | Snetkova et al, Nat Genet 2021 |
| HS2790 | chr1: 52662902-52663193 | chr1:52663061C>G | - | *COA7, TUT4* | Forebrain, Midbrain, Hindbrain, Neural Tube | Midbrain (expansion) | No | Kosicki et al, Nat Comm 2025 |
| HS2791 | chr18: 44826368- | chr18:44826694C>G | - | *SETBP1* | Forebrain Cranial | Midbrain, Hindbrain, | No | Kosicki et al, Nat Comm |
|  | 44826863 |  |  |  | nerve,<br>Hindbrain | Neural Tube |  | 2025 |
| HS2792 | chr4:<br>104425021-104425408 | chr4:104425251C>G | - | *CXXC4* | Midbrain,<br>Hindbrain,<br>Neural<br>Tube | Forebrain,<br>Face | No | Kosicki et al,<br>Nat Comm<br>2025 |
| HS1431 | chr8:<br>48581590-48583174 | 12 bp mutation | - | *SNAI2* | Limb,<br>Face | Forebrain,<br>Midbrain,<br>Hindbrain,<br>Heart | No | Kosicki et al,<br>bioRxiv 2025 |
| IRF8 32+ | chr16:<br>85984933-85985113 | Insertion of high-<br>affinity<br>motifs | Lupus | *IRF8* | Common<br>Dendritic<br>Cell<br>Progenitor | Type 2<br>Dendritic Cell | No | Ou et al, Nat<br>Immunol 2024 |
| HS2565 | chr20:<br>57155027-57159433 | 12 bp mutation | - | - | Heart<br>Ventricles | Heart Atria | No | Kosicki et al,<br>bioRxiv 2025 |
| HS3117 | chr13:<br>80275542-80277542 | chr13:80276542C>G | - | *SPRY2* | Forebrain | Midbrain,<br>Hindbrain | No | Kosicki et al,<br>Nat Comm<br>2025 |
| ZRS | chr7:<br>156790708-156793079 | chr7:156791192C>G,<br>chr7:156791218A>T,<br>chr7:156791374C>A,<br>chr7:156791382T>C,<br>chr7:156791421T>G,<br>chr7:156791483C>T,<br>chr7:156791489C>T,<br>chr7:156791491G>T,<br>chr7:156791491G>C,<br>chr7:156791491G>A,<br>chr7:156791504A>G,<br>chr7:156791515T>A,<br>chr7:156791533T>A,<br>chr7:156791550T>G,<br>chr7:156791561C>G,<br>chr7:156791708C>G,<br>chr7:156791826A>G,<br>chr7:156791493A>G,<br>chr7:156791494T>A,<br>chr7:156791488A>G,<br>chr7:156791488A>C,<br>chr7:156791830T>G,<br>chr7:156791706C>T,<br>chr7:156791602C>A,<br>chr7:156791642G>A,<br>chr7:156791856T>C,<br>2% mutagenesis,<br>ETS motif optimization | Polydactyly | *SHH* | ZPA<br>(Posterior<br>limb bud) | Anterior<br>limb bud | No | Lettice et al,<br>Hum Mol Gen<br>2003, 2008;<br>Kvon et al,<br>Cell 2020;<br>Lim et al,<br>Nature 2024 |

**Table S3.** Differential expression analysis between fluorescently sorted anterior and central cells from mZRS^WT^; hZRS^404G>A^-*mCH* hindlimb buds at E11.5.

**Table S4.** Neurodevelopmental-disorder-linked non-coding variants tested in this study.

| Variant (hg38) | dbSNP ID | Associated Disorder | Enhancer<br>(Putative Target<br>Gene(s)) | Normal<br>Enhancer<br>Activity | Reproducible<br>Changes in<br>Enhancer<br>Activity | Reference |
| --- | --- | --- | --- | --- | --- | --- |
| chr1:10736809T>G,<br>chr1:10738345C>A,<br>chr1:10738907C>T | -<br>- | Autism spectrum<br>disorder | HS2094 (*CASZ1*) | Hindbrain,<br>Neural Tube | Gain in Forebrain | Shin et al, 2024 |
| chr1:44990473C>T,<br>chr1:44990931G>T | -<br>- | Neurodevelopmental<br>disorder | HS280 (*RNF220*) | Limb | Gain in Tail | Wright et al, 2015 |
| chr6:135282407A>T,<br>chr6:135282408A>G,<br>chr6:135282410C>T | -<br>-<br>- | Autism spectrum<br>disorder | HS1351 (*MTFR2*,<br>*MYB*, *SGK1*) | Eye | - | Zhou et al, 2022 |
| chr3:182569357C>T,<br>chr3:182569692C>T | -<br>- | Autism spectrum<br>disorder | MM996 (*SOX2*,<br>*SOX2OT*, *FXR1*) | Hindbrain,<br>Neural Tube | - | Grove et al, 2019 |

**Table S5.**
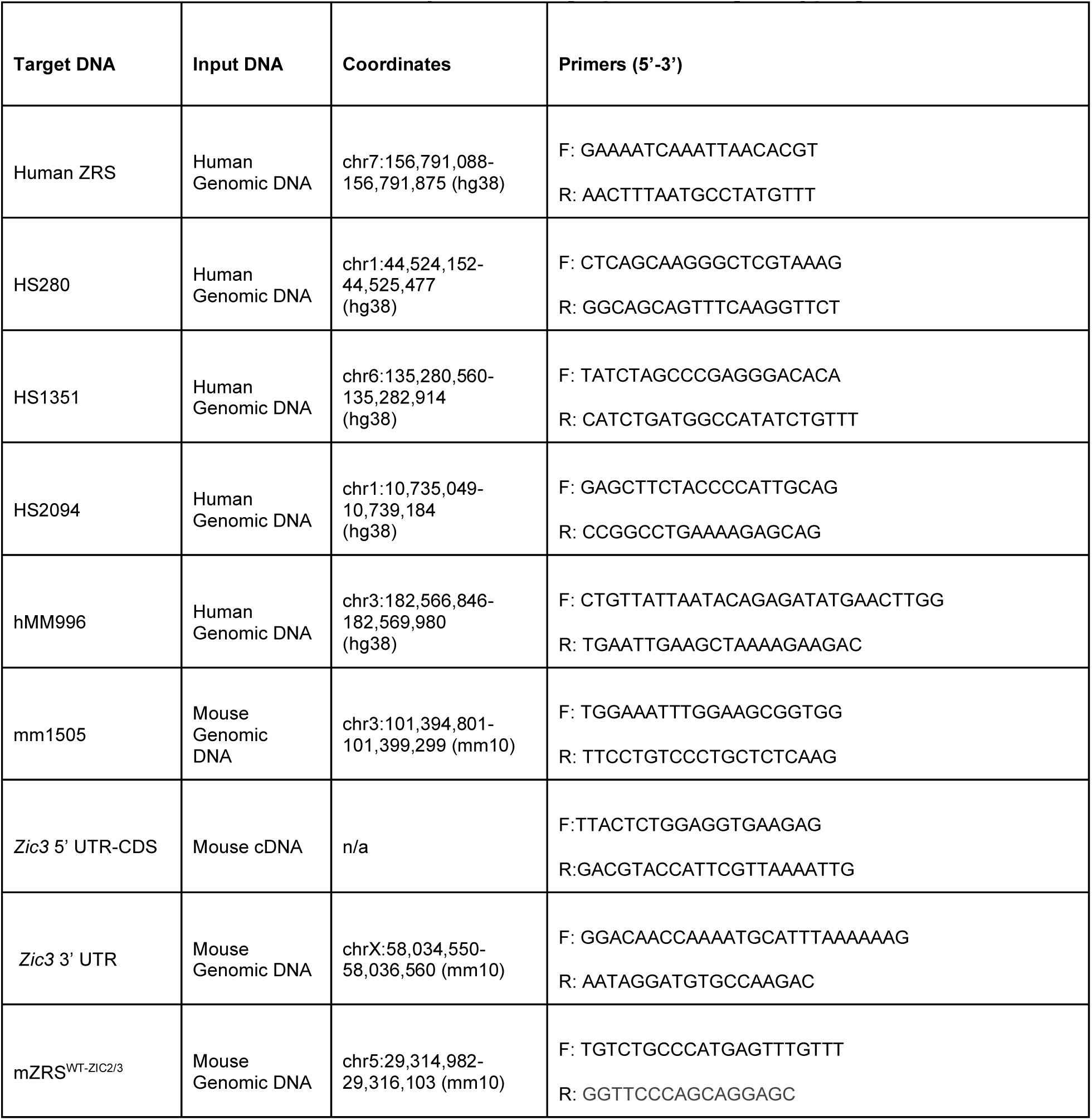
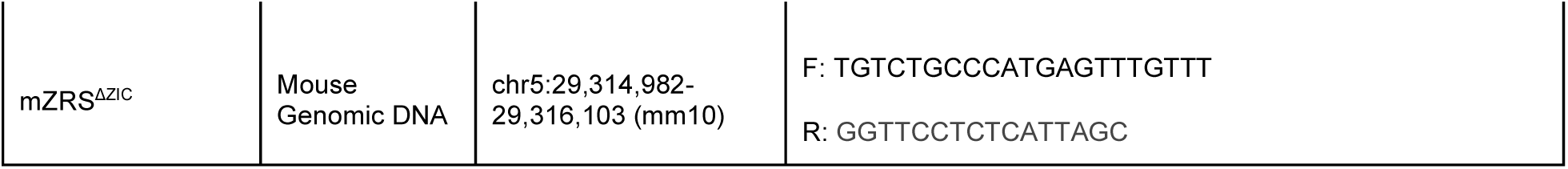
Primers used in this study for cloning, qPCR, and genotyping knock-in lines.

